# Predictive Modeling of Multiplex Chemical Phenomics for Novel Cells and Patients: Applied to Personalized Alzheimer’s Disease Drug Repurposing

**DOI:** 10.1101/2021.08.09.455708

**Authors:** Qiao Liu, Yue Qiu, Lei Xie

**Affiliations:** Department of Computer Science, Hunter College, The City University of New York; Ph.D. Program in Biology, The City University of New York; Ph.D. Program in Biochemistry, The City University of New York; Helen and Robert Appel Alzheimer’s Disease Research Institute, Feil Family Brain & Mind Research Institute, Weill Cornell Medicine, Cornell University

## Abstract

Chemical phenomics which measures multiplex chemical-induced phenotypic response of cells or patients, particularly dose-dependent transcriptomics and drug-response curves, provides new opportunities for *in silico* mechanism-driven phenotype-based drug discovery. However, state-of-the-art computational methods only focus on predicting a single phenotypic readout and are less successful in screening compounds for novel cells or individual patients. We designed a new deep learning model, MultiDCP, to enable high-throughput compound screening based on multiplex chemical phenomics for the first time, and further expand the scope of chemical phenomics to unexplored cells and patients. The novelties of MultiDCP lie in a multi-task learning framework with a novel knowledge-driven autoencoder to integrate incoherent labeled and unlabeled omics data, and a teacher-student training strategy to exploit unreliable data. MultiDCP significantly outperforms the state-of-the-art for novel cell lines. The predicted chemical transcriptomics demonstrate a stronger predictive power than noisy experimental data for downstream tasks. We applied MultiDCP to repurpose individualized drugs for Alzheimer’s disease, suggesting that MultiDCP is a potentially powerful tool for personalized medicine.

## Introduction

The target-based “one-drug-one-gene” approach has been the most dominant strategy for drug discovery and development in the past two decades, but has suffered high costs and failure rates. The phenotype-based drug discovery starts to gain increasing attention in recent years due to its ability to identify drug lead compounds in a physiologically relevant condition^1^ and the development of many high-throughput cell-based phenotypic detection methods. These technologies, including next-generation sequencing^2, 3^ and gene-editing technology such as CRISPR-Cas^4, 5^ can be applied to characterize cellular genotype and phenotype. Phenotype-based drug discovery is a target agnostic and empirical approach to exploit new drugs with no prior knowledge about the drug target or mechanism of action in a disease required^6^. The phenotype-based drug discovery methods efficiently avoid the bias on the identification of drug mechanisms of action^7^. Additionally, phenotype-based drug discovery has the power to exploit drugs for rare or poorly understood diseases^8^ such as Alzheimer’s disease.

Chemical dose-dependent response curve is a primary measure to characterize the phenotypic response of cells to the chemical treatment, for either drug sensitivity or toxicity. There are hundreds of characterized cell lines and patient samples with drug sensitivity data collected in different studies, such as Broad Institute Cancer Cell Line Encyclopedia (CCLE)^8^, Genomics of Drug Sensitivity in Cancer (GDSC)^9^, The Cancer Genome Atlas Program (TCGA)^10^. A great deal of effort has been devoted to developing machine learning models for predicting drug sensitivity^11–15^. However, the majority of previous research focused on predicting summary metrics of the drug-response curve, such as IC50 or area under the drug-response curve^16, 17^. The entire drug-response curve will provide more information than the summary metrics on the drug-response phenotype^18^. Predicting the drug-response curve is critical in the compound screening for neurogenerative diseases and autoimmune diseases, where an effective and safe drug should keep the cell alive.

The drug-response curve alone might not provide a complete picture of the drug’s actions. One rationale of phenotype-based methods is to screen and identify drugs that can modulate the global state of diseases^1^. The cellular state change could affect many different genes; thus gene expression profiles are a widely used and informative tool to characterize disease phenotype^19^. They are well suited to be used in phenotype-based drug discovery. They depict the disease state with not only the associated gene markers, but also other genes relevant to pathophysiological conditions. Moreover, the gene expression profile can characterize patients’ disease states throughout the time and give a more precise and complete diagnosis. Therefore, transcriptomics-based approach increases the capability to translate phenotype-based drug discovery to personalized medicine. Analyzing the differential gene expression profile before and after drug treatment and identifying the most regulated genes uncover the core drug mechanism of actions. This will solve the problem of drug-target deconvolution on the phenotype-based screening^20^. In the LINCS L1000 project, chemical transcriptomic data, like the differential gene expression profile perturbed by either chemicals or CRISPR-Cas technology, were collected with cost-effective and high-throughput gene expression profiling methods^21^. This project determined 978 consensus signature genes, which could reflect the cellular state pre- and post-perturbation non-redundantly and efficiently^22, 23^. It was shown that other genes expression information could be inferred from the gene expression level of these 978 genes^21^. These studies have been performed on cancer cell lines and stem cell lines collected in different tissues. Collected chemical transcriptomics data are informative for many downstream applications, including drug repurposing^24–27^, similarity-based drug candidate identification^28^, drug side effect prediction^29^, drug signature detection^30, 31^, and cellular response characterization of drug perturbations^32^.

The high-throughput methods for chemical-perturbated cell viability and transcriptome profiling have dramatically changed compound screening. However, both drug-dose response curves and chemical transcriptomics data have been collected in a very limited amount of cell lines so far and many of them do not overlap (Figure 1). For example, only ∼80 cell lines and ∼20K chemicals are tested in L1000. The exploited combinatorial space is very sparse. More importantly, enormous patient tissues (orphan diseases) are to be explored. Considering the huge combinatorial spaces of uncharacterized chemicals and cell lines or patients (dark area in Figure 1), full investigation on all the possible tests would be prohibitively expensive and time-consuming^33^. The computational approach can facilitate the drug screening process by predicting those unstudied conditions and giving informative guidance on future directions. Several state-of-the-art methods such as DeepCE and DeepCOP were designed to address the challenge for predicting chemical transcriptomics of novel chemicals for a limited number of cell lines^34, 35^.

**Figure 1.**
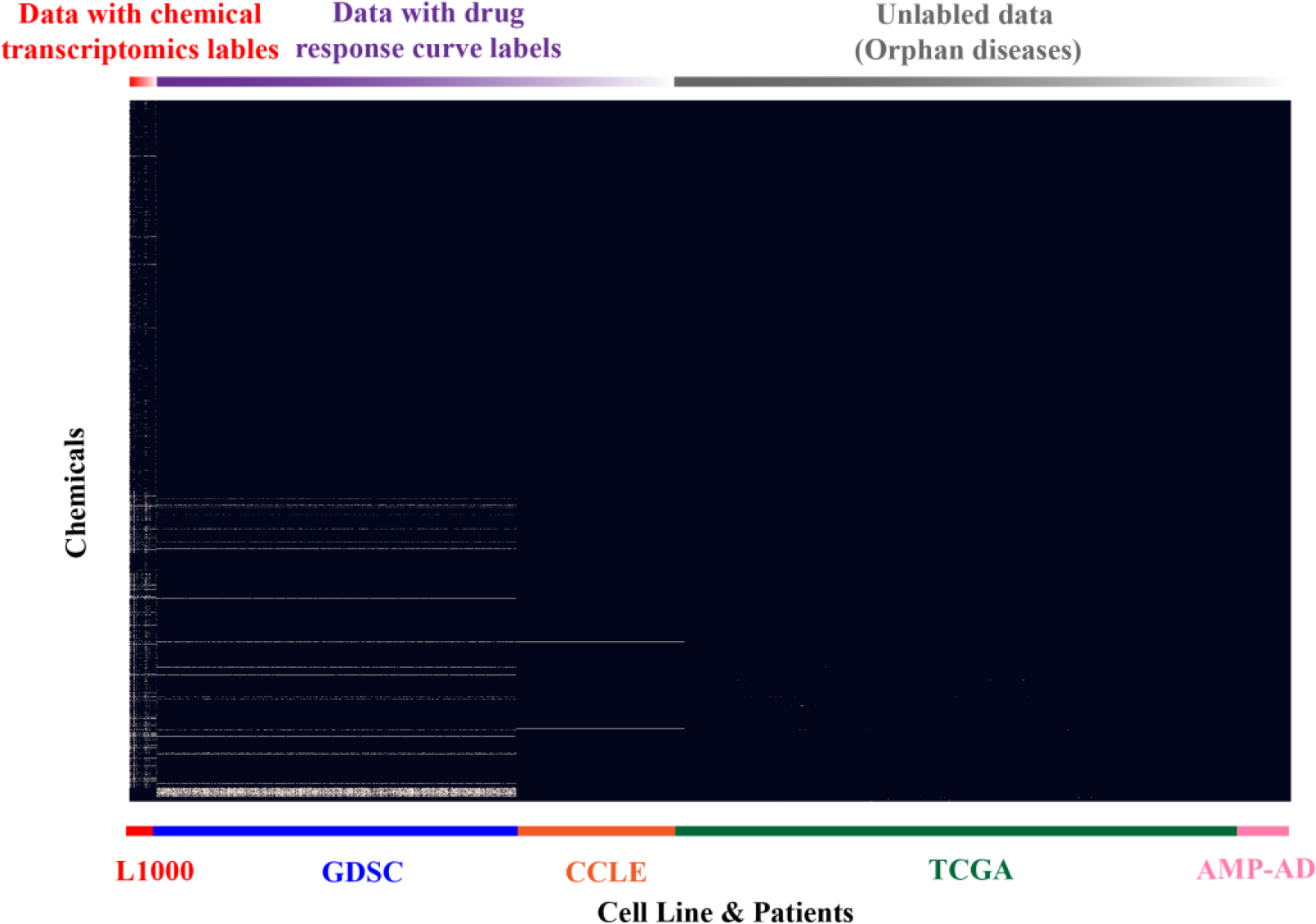
Dark chemical phenomics space explored in this study. Chemicals include all compounds in L1000, GDSC, CCLE, TCGA, and DrugBank. The cell lines/patients were collected from L1000 project, GDSC, CCLE, TCGA and AMP-AD portals. The experimentally tested drug-cell line pairs (labeled data) are marked as white dots. Noted that labeled data in L1000 and GDSC/CCLE are incoherent.

However, they still have a serious limitation on predicting chemical transcriptomics of samples that lack labels, e.g., tissues from Alzheimer’s disease patients, which are not included in L1000. Furthermore, no methods exist to predict multiplex chemical-induced phenotypic changes (chemical phenomics), including drug-response curve along with chemical transcriptomics for novel cells (or orphan diseases). Such method will no doubt accelerate the development of personalized medicines for complex diseases. Thus, we will fill this knowledge gap with our multi-task deep learning-based model in the paper.

To address unsolved challenges in the current state-of-the-art methods, we designed a new multi-task transfer learning model, MultiDCP. We show that MultiDCP significantly outperforms the state-of-the-art method DeepCE in the scenario of novel cell models. For the first time, we predict the dosage-dependent perturbed differential gene expression profile along with the drug-response curve. Furthermore, we demonstrate that the MultiDCP-predicted differential gene expression profile has stronger predictive power than the experimentally measured noisy L1000 data for a downstream side effect prediction task. This superior performance is attributed to an innovative knowledge-enabled autoencoder for gene expression profiles, integration of multiple diverse labeled and unlabeled omics data, and the joint training of the multiple prediction tasks. With the unique chemical embedding approach in DeepCE, we have the ability to explore the chemical transcriptome and drug response curve prediction problems for both novel drugs and novel cell lines or patients. These two prediction tasks are crucial for phenotype-based compound screening. We further apply MultiDCP to conduct drug repurposing for individual Alzheimer’s disease (AD) patients. The clinical potential of proposed drug leads on AD is supported by existing experimental and clinical evidence.

## Results

### Architecture of Multi-task Dose-dependent Chemical Phenomics model (MultiDCP)

The MultiDCP model integrates incoherently labeled and unlabeled data from multiple resources to perform multiple tasks including dose-dependent chemical-induced differential gene expression predictions (chemical transcriptomics) and cell viability predictions for *de novo* drugs and *de novo* cell lines (Figure 2). This model includes four input components. The first one is a graph convolutional network, extracting graphic fingerprints from chemical structures. The second component models the chemical substructure-gene interactions through an attention network^35^. The second component’s input combines the first module’s chemical graph fingerprints with the gene embedding module’s vector representations for human genes. The gene embedding module studies the gene information in a gene-gene interaction network context using a node embedding model, named node2vec ^36, 37^. The third component is a novel knowledge-enabled autoencoder module, which includes a cell line encoder and a cell line decoder. The cell line encoder compresses cell gene expression profiles to low dimensional dense vectors. Then the cell line decoder decompresses the dense vector and generates the original cell line gene expression profile from it. Different from conventional autoencoders, the cell line encoder uses a transformer module to simulate gene-gene interactions. This autoencoder module is jointly trained with other supervised learning tasks (Figure 2B). The fourth one extracts the embedding vector representation of dosage information. The outputs from these four input components are concatenated together and used as the inputs of task-specific fully connected layers for various downstream tasks, e.g., dose-dependent chemical transcriptomics prediction or cell viability prediction.

**Figure 2.**
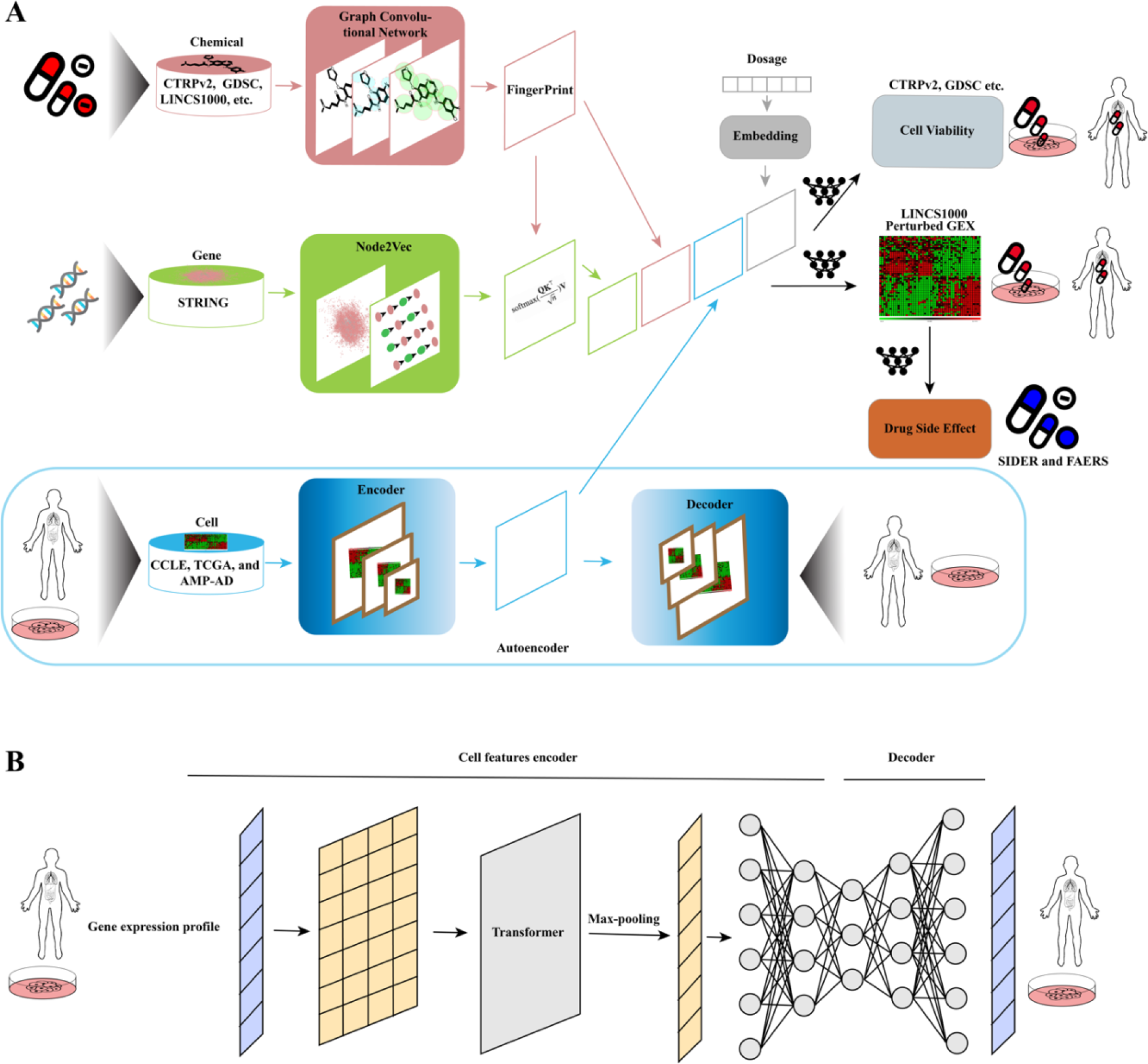
Architecture of MultiDCP and cell line autoencoder. A) The MultiDCP includes four input components: drug graphic fingerprint extraction module, gene vector extraction module, cell line encoder module and dosage embedding module. The last components are task-specific fully connected layers for various downstream tasks. B) Architecture of autoencoder. The cell line encoder will compress the gene expression profile to a low dimensional hidden vector. It includes a transformer module, a max pooling layer and fully connected layer. The decoder part will reconstruct the gene expression profile and is a fully connected module.

### MultiDCP outperforms the state-of-the-art model on predicting chemical-induced differential gene expression profiles for novel cells

The DeepCE model has shown promising results on the chemical-induced differential gene expression profile prediction task but it only focused on the novel drug setting^35^. One potential drawback of DeepCE is that it could only be applied to a limited number of cell lines due to its design in representing the cell line feature. In the input layer, it applied the one-hot encoding method to encode different cells. The first issue with this is that the model needs to modify the input dimension whenever the predictions are for new cell line types or tissues. In the inference step, because no dense vector representation has been learned in the embedding layer for new cell types or tissues, the vector has to be randomly initialized. When the number of cell types expands dramatically, the sparsity of the input will increase and this issue will become worse. Instead, we used the basal gene expression profile from a cell line or tissue as the input features. The dimension of this input is fixed with the number of consensus signature genes determined in the LINCS L1000 project. The numerical value in each dimension is the normalized basal gene expression value for each gene. Besides, this design enables us to use an autoencoder to support the learning of dense vector representation and this increases the regularization for the MultiDCP model. By adding this autoencoder, we can take advantage of unlabeled basal gene expression profile data and get a more robust vector representation. This will also increase the regularization during the learning step for MultiDCP and avoid the overfitting problem. The encoder component in MultiDCP and the cell line encoder in the autoencoder share the same parameters.

With all the designs mentioned above, we demonstrated that MultiDCP dramatically improved the model performance on the prediction of chemical transcriptomics in the novel cells setting. In this study, we applied a challenging leave-new-cells-out cross-validation strategy to evaluate the performance of MultiDCP and DeepCE. We initially separated cells into different clusters based on their cell line gene expression profiles (Supplementary Figure 1). We divided the cells into different folds in the cross-validation step and kept the cell lines belonging to the same cluster in the same fold. In other words, the cell lines in the test fold were significantly different from those in the training and development folds in terms of the gene expression profile. We compared model performance with multiple metrics, including the averages of Pearson correlation and Spearman correlation between the predicted chemical transcriptomics and the ground truth chemical transcriptomics. The Pearson correlation of MultiDCP increases by 10% in the development dataset and 15% in the test dataset compared with DeepCE. When measured by the Spearman Correlation, MultiDCP outperformed DeepCE by 15% in the development dataset and 17% in the test dataset, respectively (Figure 3A). We ranked the regulated genes based on their normalized differential expression values to top-K upregulated genes or downregulated genes. When the performance is evaluated by the top-K precision, MultiDCP also demonstrated significant performance gain over DeepCE. The predicted precision of the top-10 upregulated genes and top-10 downregulated genes increases by 19%-20% for the test dataset and 10%-13.5% for the development dataset (Figure 3B and 3C). The predicted precision of top-100 upregulated genes and top-100 downregulated genes increases by 12%-18% for the test dataset and 7%-11% for the development dataset (Figure 3B and 3C).

**Figure 3.**
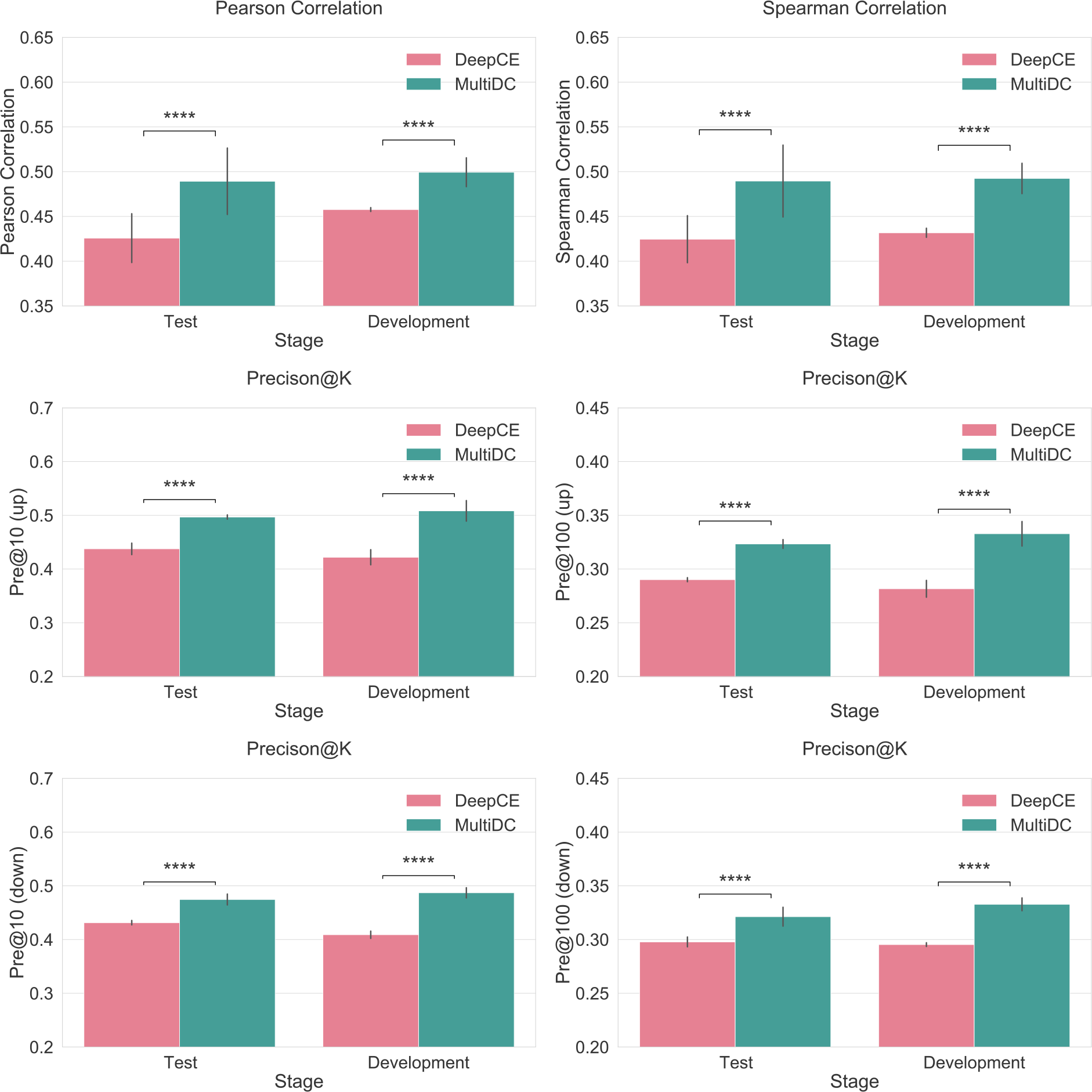
Comparison of drug perturbed gene expression profile prediction performance of DeepCE and MultiDCP. A) The average Pearson correlation and Spearman correlation of predicted gene expression profile and ground truth are used as the evaluation metrics. B) Comparison of drug perturbed gene expression profile prediction precision of the top upregulated genes (Pre@K (up)) and top downregulated genes (Pre@K (down)) of DeepCE and MultiDCP. We selected the top 10 and 100 upregulated genes and downregulated genes and evaluated the prediction precisions for them. **** indicates that the p-value of two-tailed t-test is less than 1.0x10^-4^.

### Joint unsupervised and supervised training improves the performance of MultiDCP

One of the critical innovations in MultiDCP is the knowledge-enabled autoencoder which supports the learning of vector representation of cells and patients (Figure 2B). In the benchmark studies, the autoencoder component was trained with heterogeneous data from two widely used cancer-related databases, CCLE^26^ and TCGA^8, 10^. Around 1.3K cancer cell lines and 10K patient tissue gene expression profiles were extracted from these two databases followed by the removal of the batch effects. The autoencoder’s input and output are the expression values of the 978 consensus signature genes determined in the LINCS L1000 study, which represent the phenotypic response induced by drugs. We also tested other gene-specific feature profiles as the input of the autoencoder. However, we showed that the models with gene dependency profiles as cell features had inferior performance than using gene expression profiles (Supplementary Table 1). We tested two different strategies to train the autoencoder in MultiDCP, pre-training-fine-tuning, and jointly-training. In the pre-training-fine-tuning strategy, we first trained the autoencoder model with the gene expression profile data completely. Next, we initialized the cell features encoder in MultiDCP with the parameters learned in the pre-training stage for the cell line encoder of the autoencoder. We then fine-tuned them together with other parameters with the downstream supervised learning tasks. Within the jointly training strategy, we coupled the training of those two aforementioned encoder components together and trained autoencoder and MultiDCP in an alternative way. During the training stage, the parameters for the encoder part in both models were shared and updated simultaneously. For both learning strategies, we utilized the leave-new-cells-out cross-validation to test the model performance. We carefully split the train-development-test dataset so that the cells in the training dataset for the autoencoder were consistently kept for the supervised MultiDCP training. The same criteria were applied to the development and test dataset (Supplementary Figure 2). We found that the jointly training strategy showed superior performance to the pre-training-fine-tuning strategy (Table 1). The jointly training procedure could increase performance by 7%-15% for Pearson Correlation and 7%-15% for Spearman Correlation, respectively. Besides, we also compared the prediction precision for the top K upregulated and downregulated genes (Table 1). The results agree with the conclusions from the other metrics. We believed that joint training could provide more regularizations on both tasks.

**Table 1.**
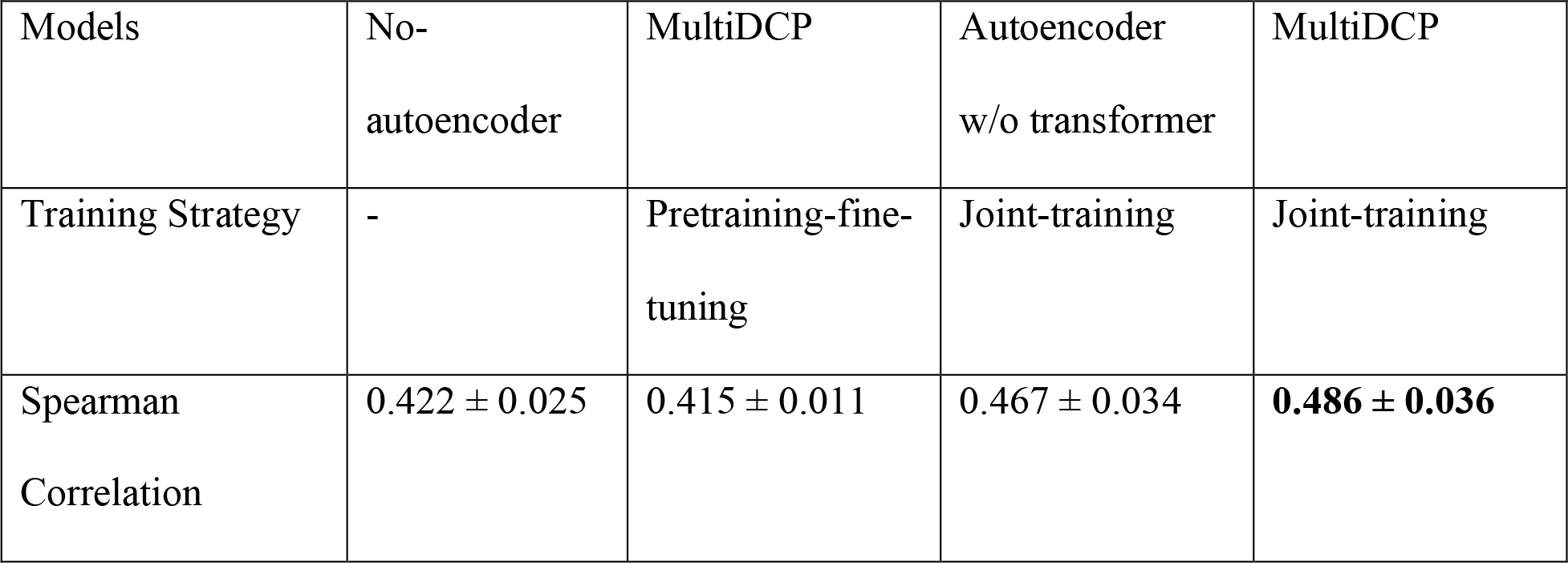
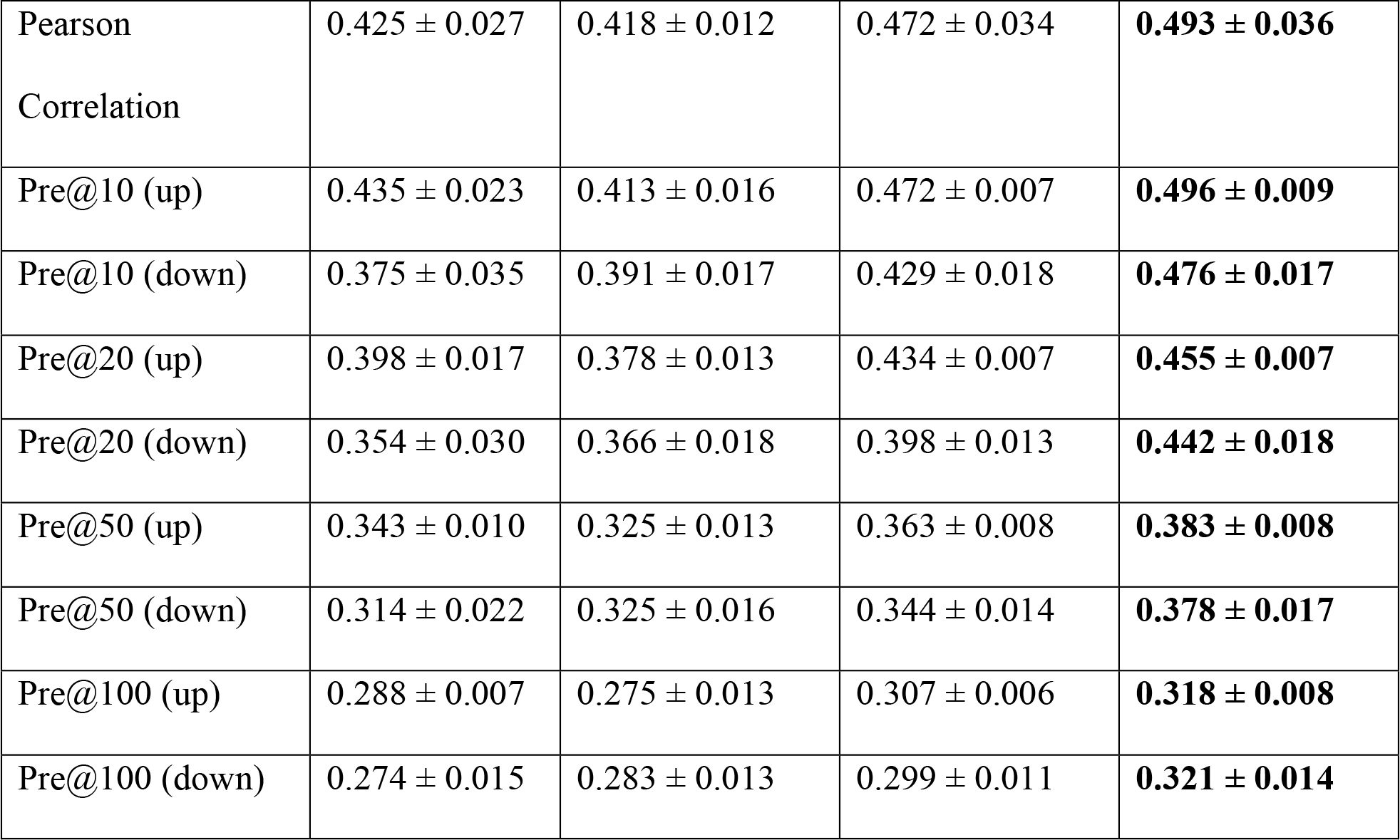
The results of the ablation test. The no-autoencoder training strategy indicates that neither the model is trained together with the autoencoder model nor the encoder parameters are initialized with the pretraining model. The pretraining-fine-tuning strategy means that the cell features encoder parameters are initialized with the pretraining model. The jointly training strategy means that the autoencoder and supervised learning components are couples and jointly trained. We selected the Pearson correlation and Spearman correlation as the comparison metrics. We also selected the top 10, 20, 50, 100 upregulated and downregulated genes and evaluated the prediction precisions of them.

### Attention-based autoencoder improves the performance of MultiDCP

Another innovative component in the encoder module is a transformer component. The transformer uses a self-attention mechanism to model gene-gene interactions. The transformer module has been shown to successfully boost model performance in many applications and different areas, such as Natural Language Processing, Computer Vision, biological sequence modeling, and drug discovery applications^38–43^. We performed an ablation study to test the importance of this transformer. We deployed another baseline model which has a similar structure as vanilla autoencoder (autoencoder w/o transformer). To make an apple-to-apple comparison, we kept all the other components the same, including the decoder part in the autoencoder and the other components in the MultiDCP model, except that this baseline model doesn’t have the transformer module. We demonstrated that the transformer module was critical for the model to have superior performance for all the metrics we measured. Specifically, the Pearson correlation and Spearman correlation of the transformer-enhanced autoencoder increased by 4%-5% compared with the baseline model. The prediction precision for the top 10 upregulated genes and downregulated genes increased by 5%-10%. The prediction precision for the top 100 upregulated genes and downregulated genes increased by 4%-7% (Table 1).

### Teacher-student training improves the model performance

We were inspired by the idea of the teacher-student model and implemented an innovative way to improve the model performance because the training dataset is limited. Based on the average Pearson correlation score among bio-replicates, our data were separated into high-quality datasets and low-quality datasets. The samples with score higher than 0.7 were defined as high-quality data. Then we augmented data with the procedure we described in Methods. We repeated the data augmentation loop four times. During each loop, we included additional data samples with their predicted gene expression profile. It demonstrates that the model trained with augmented data can outperform the model trained without it by 5-7% (Figure 4). It is also worth noting that this result is gathered in the scenario of leaving new cells out.

**Figure 4.**
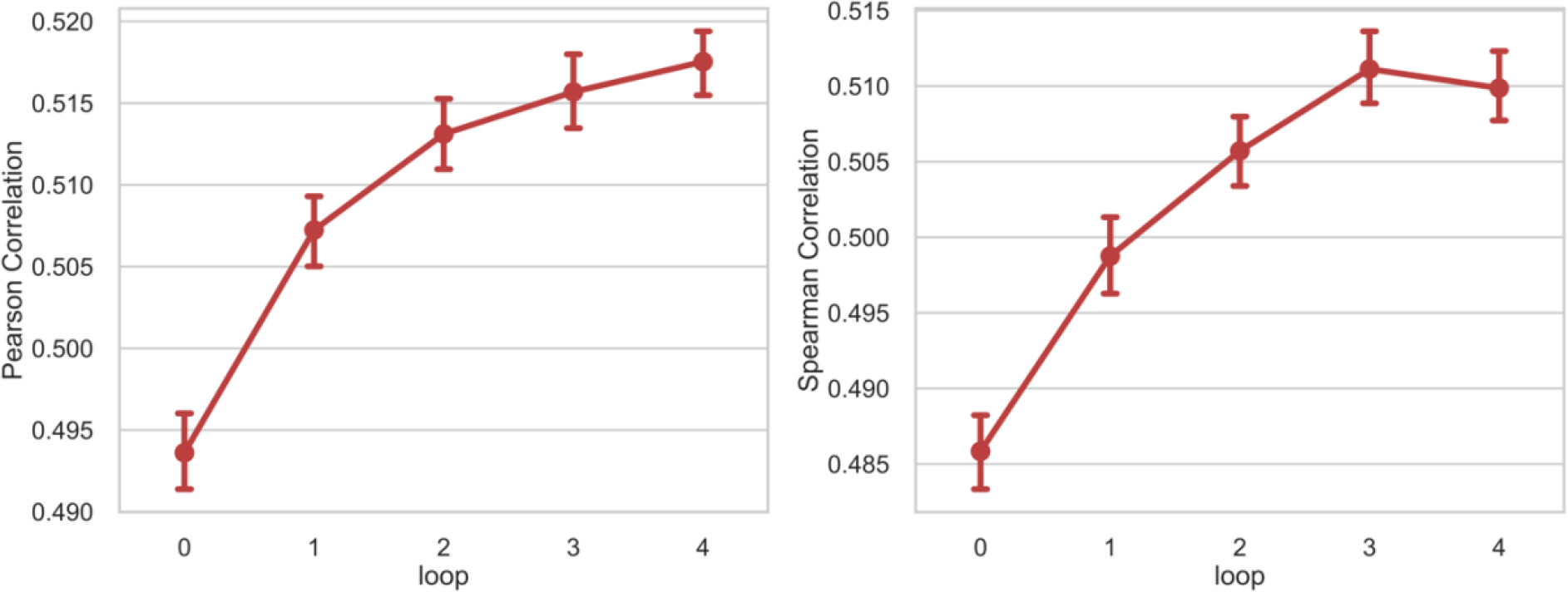
The changings of the Pearson correlation and Spearman correlation during the process of data augmentation. Following the approach outlined in Methods, we added extra data after each loop of data augmentation. Loop 0 means that we did not add any data with predicted gene expression profile.

### MultiDCP shows potential in predicting chemical dosage-dependent curve of cell viability

The drug response curve shows whether drugs are effective or safe in treating diseases. More importantly, it can indicate the optimal drug dosage to use. Such information cannot be obtained from IC50 or area under the drug response curve that are the two most widely used summary metrics for drug sensitivity prediction. For the first time, we could use MultiDCP model to predict dosage-dependent cell viability at multiple dosages. It is worth mentioning that many published models for drug sensitivity prediction usually ignore the dosage information^16, 17, 44^. Because the collected drug response data are heterogenous from different resources, the range of dosages used in each study is not consistent ^45, 46^. To solve this problem, we first fit the heterogenous drug response data to drug response curves with the procedure mentioned in Methods. These drug response curves were used to extract cell viability at selected dosages so that we could have drug response data at the same dosage range across different databases. The MultiDCP model showed promising results on the prediction of drug response curves. Figure 5 shows two examples of predicted drug response curves. Overall, the Pearson correlation and Spearman correlation between predicted drug response curves and ground truths are 0.802 and 0.782, respectively (Table 2). We also evaluated the drug-wise and cell-wise Pearson correlation and Spearman correlation by averaging the metrics for each drug and each cell, respectively (Table 2).

**Figure 5.**
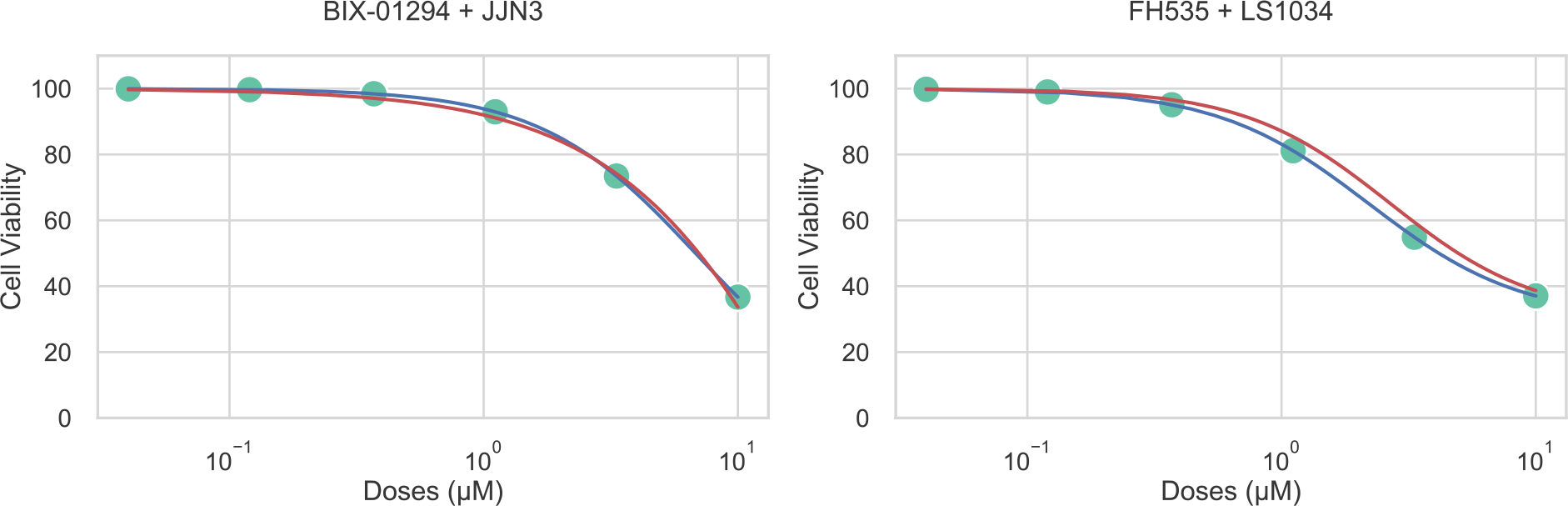
Comparison of predicted drug response curve (red) vs ground truth drug response curve (blue) for two examples, BIX-01294 and JJN3 pair and FH535 and LS1034 pair.

**Table 2.** The performance of MultiDCP cell viability model (MultiDCP-CV) on cell viability prediction. The Pearson correlation and Spearman correlation are used as the evaluation metrics.

|  | Pearson Correlation | Spearman Correlation |
| --- | --- | --- |
| Point wise | $0.803 \pm 0.013$ | $0.780 \pm 0.015$ |
| Drugs wise | $0.721 \pm 0.068$ | $0.632 \pm 0.052$ |
| Cells wise | $0.766 \pm 0.087$ | $0.687 \pm 0.103$ |

### Predicted chemical transcriptomics profile has strong predictive power for downstream task

L1000 chemical transcriptomic profiles are excellent features for predicting organismal phenotypes such as drug side effects^47^. However, many existing drug-induced differential gene expression profiles are noisy due to the existence of various confounding factors in experiments^35^. These noisy chemical transcriptomic data are not reliable for other downstream prediction tasks. Many works have focused on improving the data quality by removing these noisy data or developing new deconvoluting methods for data processing^23, 48^. We designed an experiment to test whether our predicted chemical transcriptomics could be more effective on the downstream learning tasks than the unreliable experimental chemical transcriptomics data. We focused on the drug side effect prediction that only uses the drug perturbed gene expression profile as input. Drug side effects were categorized to different Preferred Terms (PTs) coded in Medical Dictionary for Regulatory Activities (MedDRA) v16.0^49^. We collected data from two adverse drug reaction (ADR) datasets, an on-label ADRs side effect resource (SIDER) and an off-label ADRs PharmGKB Offsides table from FDA Adverse Event Report System (FAERS)^50, 51^. We compared the performances of ADR prediction on both datasets using both the experiment-determined noisy L1000 level-5 data and predicted perturbed gene expression profile (Table 3 and 4). We selected four classification models that have been used in the drug side effect prediction task, including logistic regression (LR), Extra Trees (ET), Random Forest (RF), and a deep neural network (DNN). All of these models were used in a multi-label classification setting and evaluated with a 5-fold cross-validation method. For the SIDER dataset^50^, the macro-ROCAUC was 7%-10% higher when we trained the models with predicted chemical transcriptomics rather than experiment-determined noisy data. For FAERS dataset^51^, we observed the same pattern. The macro-ROCAUC was improved by 3%-4%. The performances of the micro-ROCAUC have similar trends for both datasets. Considering that this dataset is imbalanced and the number of positive samples in each category are very different, the macro-ROCAUC is a more suitable metric for the performance comparison. This indicates that the predicted chemical transcriptomics are more informative than the unreliable experimental data, such as for the downstream task of side effect prediction.

**Table 3.** Comparison of the performance of drug side effect prediction models, Random Forest, Extra Trees, Logistic regression and Neural Network when using different input drug perturbed differential gene expression profile features on two different datasets, SIDER and FAERS. The evaluation metrics are macro-ROC-AUC and micro-ROC-AUC. In the input column, the predicted DGX means that the input feature is predicted by MultiDCP model and the experimental DGX means that the input feature is the profile collected by experiment but only the unreliable ones are used.

| Dataset | Input | Random Forest |  | Extra Trees |  | Logistic regression |  | Neural Network |  |
| --- | --- | --- | --- | --- | --- | --- | --- | --- | --- |
|  |  | macro | micro | macro | micro | macro | micro | macro | micro |
| SIDER | Experimental | 0.502 ± | 0.734 ± | 0.502 ± | **0.736 ±** | 0.503 ± | 0.743 ± | 0.503 ± | 0.731 ± |
|  | DGX | 0.021 | 0.023 | 0.021 | **0.025** | 0.022 | 0.024 | 0.022 | 0.023 |
|  | Predicted | **0.522 ±** | **0.742 ±** | **0.525 ±** | **0.736 ±** | **0.520 ±** | **0.747 ±** | **0.539 ±** | **0.733 ±** |
|  | DGX | **0.022** | **0.024** | **0.019** | **0.020** | **0.022** | **0.022** | **0.024** | **0.026** |
|  | p-value of t-test | 0.010 | 0.186 | 0.002 | >0.5 | 0.029 | 0.328 | <0.001 | 0.419 |
| FAERS | Experimental | 0.503 ± | **0.752 ±** | 0.509 ± | 0.739 ± | 0.505 ± | **0.754 ±** | 0.518 ± | 0.749 ± |
|  | DGX | 0.017 | **0.019** | 0.016 | 0.017 | 0.015 | **0.016** | 0.015 | 0.017 |
| | Predicted | **0.550 ±** | 0.751 ± | **0.547 ±** | **0.745 ±** | **0.554 ±** | 0.745 ± | **0.564 ±** | **0.751 ±** |
|  | DGX | **0.016** | 0.018 | **0.016** | **0.017** | **0.016** | 0.018 | **0.015** | **0.018** |
|  | p-value of t-test | <0.001 | 0.445 | <0.001 | 0.194 | <0.001 | 0.097 | <0.001 | 0.386 |

**Table 4.** Comparison of the performance of drug side effect prediction models, Random Forest, Extra Trees, Logistic regression and Neural Network when using different input drug perturbed differential gene expression profile features on two different datasets, SIDER and FAERS. The evaluation metrics are macro-PR-AUC and micro-PR-AUC. In the input column, the predicted DGX means that the input feature is predicted by MultiDCP model and the experimental DGX means that the input feature is the profile collected by experiment but only the unreliable ones are used.

| Dataset | Input | Random Forest |  | Extra Trees |  | Logistic regression |  | Neural Network |  |
| --- | --- | --- | --- | --- | --- | --- | --- | --- | --- |
|  |  | macro | micro | macro | micro | macro | micro | macro | micro |
| SIDER | Experimental | 0.362 ± | 0.364 ± | 0.393 ± | 0.291 ± | 0.363 ± | 0.343 ± | 0.353 ± | 0.331 ± |
|  | DGX | 0.017 | 0.021 | 0.016 | 0.013 | 0.018 | 0.021 | 0.022 | 0.023 |
|  | Predicted | **0.412 ±** | **0.382 ±** | **0.436 ±** | **0.371 ±** | **0.375 ±** | **0.374 ±** | **0.384 ±** | **0.394 ±** |
|  | DGX | **0.024** | **0.018** | **0.029** | **0.019** | **0.024** | **0.022** | **0.024** | **0.026** |
|  | p-value of t-test | <0.001 | 0.002 | <0.001 | <0.001 | 0.029 | <0.001 | <0.001 | <0.001 |
| FAERS | Experimental | 0.101 ± | 0.222 ± | 0.097 ± | 0.224 ± | 0.095 ± | 0.214 ± | 0.103 ± | 0.214 ± |
|  | DGX | 0.015 | 0.013 | 0.011 | 0.015 | 0.013 | 0.013 | 0.015 | 0.017 |
|  | Predicted | **0.251 ±** | **0.252 ±** | **0.258 ±** | **0.271 ±** | **0.255 ±** | **0.251 ±** | **0.258 ±** | **0.250 ±** |
|  | DGX | **0.013** | **0.016** | **0.013** | **0.015** | **0.016** | **0.014** | **0.015** | **0.018** |
|  | p-value of t-test | <0.001 | <0.001 | <0.001 | <0.001 | <0.001 | <0.001 | <0.001 | <0.001 |

### Personalized Drug Repurposing for Alzheimer’s Disease

Drug response data collected on Alzheimer’s Disease (AD) patient tissues are extremely rare. Thus, predicting potent treatment for AD patient is a very challenging task using the majority of existing methods that are incapable of predicting chemical phenomics for novel cells or tissue. We applied MultiDCP to this challenging task and the results were promising. 46 AD patients from AMP-AD project were selected as described in the Method. We predicted perturbed differential gene expression profile induced by drugs accumulated in DrugBank^52^ for these 46 AD patients. Each patient’s predicted perturbed gene expression profile by MultiDCP and the up-regulated and down-regulated gene sets when compared with healthy controls were used to discover the potential drug candidate as AD therapy by performing GSEA analysis^53^. We aggregated the top-ranked candidate drugs and performed drug-set enrichment analysis based on DrugBank category terms. We then selected the enriched terms with p-value < 10^-3^ based on hypergeometric test results (Supplementary Table 2). The drug categories related to neural system, such as neural transmitter agents, were significantly enriched. Among the drug candidates that we identified, Acetylsalicylic acid and Mefenamic acid are anti-inflammatory drugs that are also related to the neural system (Supplementary Table 3 and 4). Mefenamic acid has been proven to have neuroprotective effects and improves cognitive impairment in *in vitro* and *in vivo* Alzheimer’s disease models^54^. Simiarly, acetylsalicylic acid, as a nonsteroidal anti-inflammatory drug, also showed protential in treating AD^55^. Among the targets of neural system related drugs, GABA receptors and dopamine receptors were known to play roles in AD^56, 57^. β-adrenergic receptors, histamine receptors, 5-HT receptors and adenosine receptors had been studied as potential therapeutic targets of AD ^58–62^. For example, pentazocine and cilostazol showed anti-inflammatory effects^63^ and preventive effects on cognitive decline in AD patients^64^, respectively. Furthermore, MultiDCP predicted the drug response curve given a drug and a AD patient expression profile (Figure 6, Supplementary Figure 3, 4 and 5). We prioritized the drugs which won’t cause toxicity on the patient tissue, as defined as the cell viability larger than 90% at the drug concentration of 1 µM. Among the four aformentioned drug candidates, only pentazocine shows toxicity at 1 µM (Figure 6).

**Figure 6.**
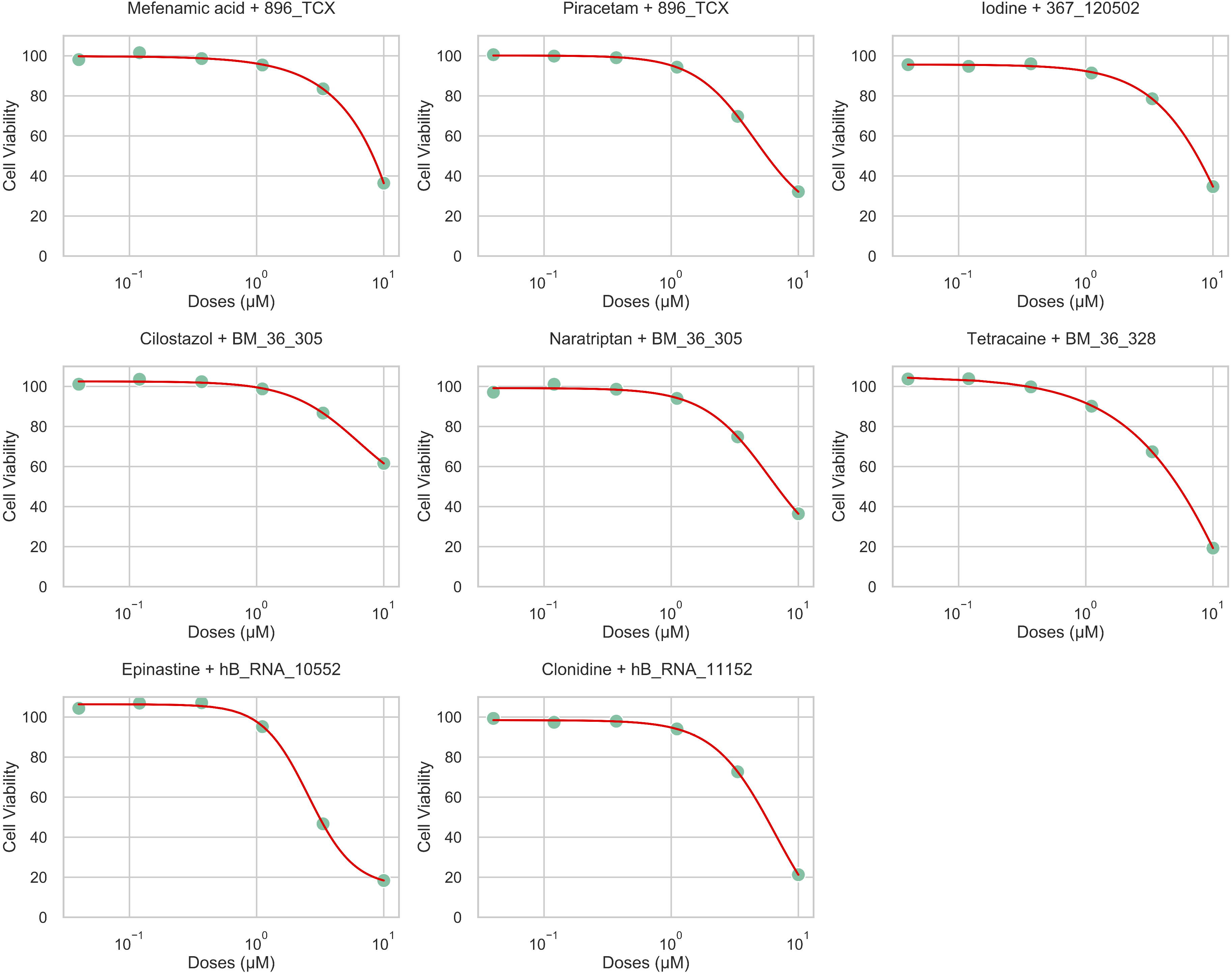
Predicted drug response curves for selected drug and AD patient pairs. The pentazocine shows a high toxicity at 1 µM for AD patient BM-36-383, while the others show mild effect for their corresponding patients.

We further clustered the AD patients based on their basal gene expression profiles. 46 AD patients were grouped into 3 clusters, each of which have 29, 8 and 9 patients. To characterize the patient subgroup, we summarized the disease signature for each subgroup by using all genes that are up- or down-regulated in more than one patient. We then perfomed functional annotation clustering with DAVID ^65^ for the disease signature (Supplementary Table 5). We found cyclin, that controls the modulation of neurotransmitter, steriod, and cell cycle related pathways are enriched in genes from subgroup 1, subgroup 2, and subgroup 3, respectively. Previous research has shown the dysregulation of cyclin and steroid is related to AD^66, 67^. The re-entry of cell cycle in neuronal cell could be the cause of AD^68^. Although the drug category related to nervous system are enriched in all 3 clusters, the drug category “neural transmitter agent” is more frequent in cluster 1, while the terms like “steroids” and “hormone” appears more frequent in cluster 2 and 3 (Supplementary Figure 6). Thus, results from drug set enrichment analysis and patient based GSEA analysis are consistent. These results indicate that our framework could be used for personalized AD drug repurposing.

## Discussion

It is nearly impossible to fully explore the combinatorial space of novel drugs and novel cell lines or patients experimentally. Thus it is crucial to develop a computational approach to uncover this chemical phenomics space^33^. In this paper, we showed that our model, MultiDCP, filled in many gaps and outperformed the state-of-the-art model on the chemical phenomics prediction for novel cells, including cultivated cell lines and patient tissues. Combined with the novel chemical representation module in DeepCE, our model can satisfy the prediction of chemical phenomics under the circumstance of both novel cells/patients and novel drugs^35^. We demonstrated that the predicted chemical transcriptomics were informative to downstream applications. The core component of MultiDCP on solving the novel cell problem is unsupervised representation learning with heterogeneous data sets. The core module for the cell representation can be translated and applied to other drug discovery applications. Another novelty of our model is that our prediction is dosage dependent. To demonstrate how useful our model’s predictions are on downstream application, we conducted a comparison of our predicted data and experimental unreliable data for the drug side-effect prediction, which showed that our predicted data is more informative than the experimental unreliable data. Our model is designed to utilize the patient gene expression profile naturally, so we believe that our model can be easily expanded for personalized medicine.

Many practitioners are concerned about applying phenotype-based drug discovery strategies because the drugs discovered with this strategy lack information about the mechanisms of action in the endpoint^6^. The breakthrough of this problem is the emergence of the high throughput genetical and molecular technology. These techniques help delineate the cellular models and diseases. The gene expression profile is one of the widely used strategies on phenotyping cellular models with the advent of sequencing technology. With the help of a system biology approach, the core mechanism behind the scenes could be uncovered more effectively. Many system biology works have illustrated that combining the pathways and network information with molecular phenotyping information could help to determine the mechanism of action^69^. Our predicted perturbed gene expression profile, as the primary resource in this scenario, will have broad applications. It will help phenotype-based drug discovery overcome the aforementioned limitations on understanding drug mode of actions, while still taking advantage of the exploitation power on discovering novel first-in-class drugs with cellular assays or even some poorly investigated disease models. Further development of MultiDCP will make it possible to perform personalized phenotype compound screening using patient data directly.

## Methods

### Data set

#### Bayesian-based peak deconvoluted LINCS L1000 dataset

The LINCS L1000 dataset provides comprehensive perturbagen induced differential gene expression profiles for 50K perturbagens and around 80 cell lines. The perturbagens include drugs, RNAis, and CRISPR-Cas assays. We used the gene expression profile that are inferred with a more robust Bayesian-based peak deconvoluted approach^23^. Each sample is a unique drug, cell line, dosage, and time combination and the profile include the differential gene expression of the 978 landmark genes determined in the LINCS L1000 project. We used the precomputed level 5 data available at https://github.com/njpipeorgan/L1000-bayesian. Only the high quality and reliable data are selected following the same procedure in a recent study. It was shown that the data quality would affect the prediction performance^35^. To define the reliability, we firstly found all biological replicates for a sample and computed the average Pearson’s correlation score among replicates. The samples with the average Pearson’s correlation score larger than 0.7 are defined to be the high-quality data. The samples with the average Pearson’s correlation score less than 0.7 were not included for model training purposes but were used in the following data augmentation steps.

#### Cell gene expression profile input dataset

We use the heterogeneous basal gene expression profiles collected in CCLE and TCGA, which include around 13K cell lines and 11K patient tissue samples, respectively. Different from the characteristic direction (CD) method used by LINCS L1000 technology, the gene expression data were collected with RNAseq technology in CCLE and TCGA program^8, 10^. To reduce the error introduced by different technologies, we only used the cell gene expression profile collected with RNAseq technology. First, we integrated the RNAseq raw counts data from all databases and removed the batch effects with the Combat_seq function in the sva package^70^. The final input is log2 transformed with a pseudo-count of 0.001 (Supplementary Figure 7). 15 cell lines were found to exist in both the LINCS L1000 dataset and the collected RNAseq dataset.

#### Cell viability prediction dataset

The cell viability dataset was retrieved from the integrated drug response database PharmacoDB^71^. They integrated the data from the CCLE dataset, GDSC1000 dataset, gCSI dataset, FIMM dataset, and CTRPv2 dataset. Only cell lines, which could be found in CCLE, are used because we need the cell line gene expression profile as the model input. The number of common cell lines from each different dataset with CCLE dataset are shown in (Supplementary Figure 8). Since we just need the drugs SMILES as the input for drugs, all the drugs are kept in the dataset. We fit the drug response curve with the following equation (1):

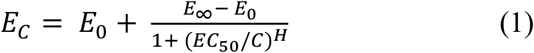

where the *E*_0_ is initial cell viability, *E*_∞_ is the infinity cell viability, *EC*_50_ is the dosage at which the cell viability is 50%, *C* is the dosage, and *H* is the hill plot slope. The data that are not able to fit Equation (1) are removed. For example, some data have *EC*_50_ value as infinity or not determined. We then calculated the cell viability *EC* at the desired dosage range. We noticed that the cell viability did not change in the desired dosage range in some samples. In this scenario, we only used the data at the maximum dosage and minimum dosage. This strategy removes many noisy data points that can be seen from the distribution before and after the removal (Supplementary Figure 9). After filtering, the dataset has 373 drugs and 886 cell lines (Supplementary Table 6).

#### Drug side effect prediction dataset

We used two adverse drug response datasets in the side effect prediction study (Supplementary Table 7). The on-label adverse drug responses side effect resource (SIDER) dataset has 834 marketed drugs, 3,166 adverse drug response preferred terms, and 88,635 drug-ADR associations^50^. The off-label ADRs PharmGKB Offsides table from FDA adverse event report system (FAERS) has 684 drugs, 9,405 ADR terms, and 26,0238 drug-ADR associations^51^. For the ADR terms, we also removed the ADR terms that had only less than 10 drugs. The ADR terms used in SIDER and FAERS were labeled with the preferred terms from MedDRA v16.0^49^.

### MultiDCP architecture

#### Overall architecture

The MultiDCP architecture concatenates the hidden representations from four types of inputs, drugs, cells, genes, and dosages. The drugs are inputted as SMILE strings. The SMILE strings are used to extract the chemical structure information with RDKit^72^, including atom and bond information. The atom and bond information are inputted into a graph convolutional network (GCN) layer to output the graphic fingerprint for the chemicals. In the LINCS L1000 high-quality dataset, 527 drugs’ perturbed gene expression profiles were collected for the initial training. The cells are represented with the gene expression profile and transformed to low dimensional hidden representations through a transformer boosted encoder. 15 cell lines were included in the dataset from LINCS L1000. The autoencoder, which shares the same parameter with this encoder, was jointly trained with gene expression profile collected from CCLE and TCGA. We used the STRING protein-protein interaction dataset for gene embedding purposes, which includes 19K genes and 12M interactions. Gene embedding vectors for the 978 landmark genes were learned with the node2vec algorithm^36^. The dosage was encoded with one-hot encoding method and its hidden representation was extracted from an embedding layer.

#### Graphic Neural fingerprint

GCN has shown to be an efficient way of extracting 2D chemical structure information^35^. Thus, we applied a GCN module to get the graphic neural fingerprint of chemicals with the following equation (2):

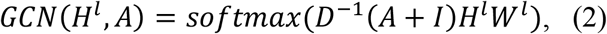

where the *H^l^* is the hidden representation in layer *l*, *A* is the graphic adjacency matrix, *I* is an identity matrix, *D* is the node degrees diagonal matrix for (*A*+*I*) and *W^l^* is GCN model parameters in layer *l*. The nodes of the chemical structure graph are atoms, and the edges of the graph are chemical bonds. The atoms information and bonds information were extracted from the chemical SMILES strings with RDKit^72^. Intuitively, the multi-layer graph convolutional layers can represent chemical substructures that are centered at the granularity of each atom and composed of neighbor atoms within a multi-hops distance.

#### Autoencoder

The cell gene expression profile autoencoder was trained with data from CCLE and TCGA. It includes two components, an encoder, and a decoder (Figure 2B). The encoder includes a gaussian noise and dropout layer, a transformer module, and a max-pooling layer. The gaussian noise and dropout layer was used to introduce noise into the input during the training process. It is worth mentioning that these two modules would not be used during the inference process. The transformer stage was used to extract the gene-gene interaction information. The transformer itself also includes an encoder component and a decoder component. Both encoder and decoder include attention-based sublayers, which have attention modules, add & norm stages, and feed-forward stages.

### Model training

#### Training procedure

We applied two different strategies on the model training of MultiDCP. MultiDCP have two different major modules, the autoencoder and the downstream task prediction module. When training with pretraining and fine-tuning procedure, we firstly train the autoencoder module with heterogeneous cell line profile datasets (Procedure 1). The parameters in the encoder part will be saved. During the fine-tuning stage, the encoder part of downstream task prediction module is initialized with the saved parameters from the autoencoder. Then the parameters in the downstream task module will be tuned with specific supervised learning task. We also utilized a jointly training procedure (Procedure 2). In this procedure, we trained the autoencoder and downstream task in alternative way. It worth mentioning that the encoder is shared for the autoencoder and downstream task. In each epoch, we firstly updated the parameters of the autoencoder with the reconstructive loss, then updated the parameters of the downstream tasks.

##### Procedure 1 Pretraining-fine-tuning procedure

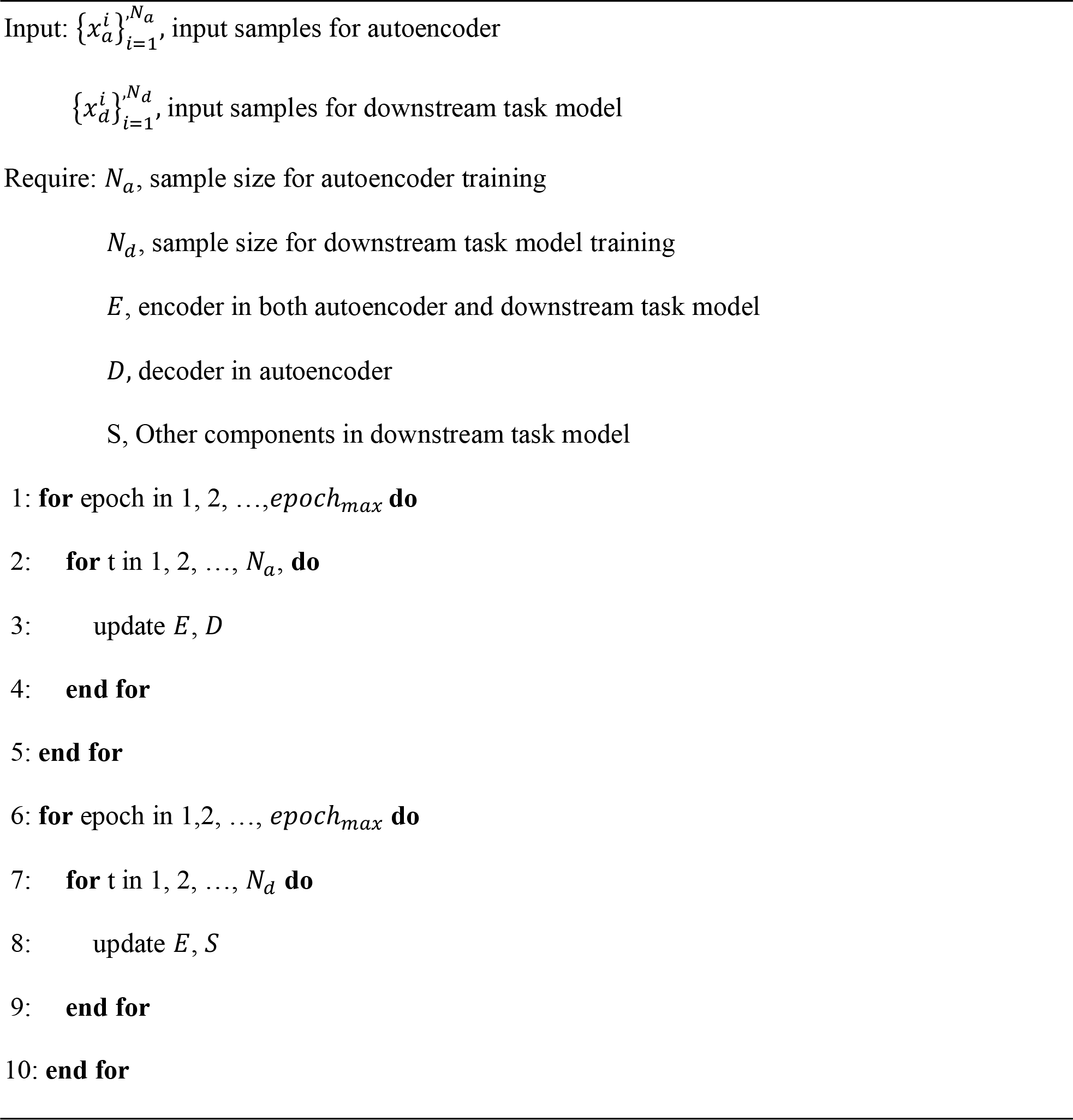

##### Procedure 2 Pretraining-fine-tuning procedure

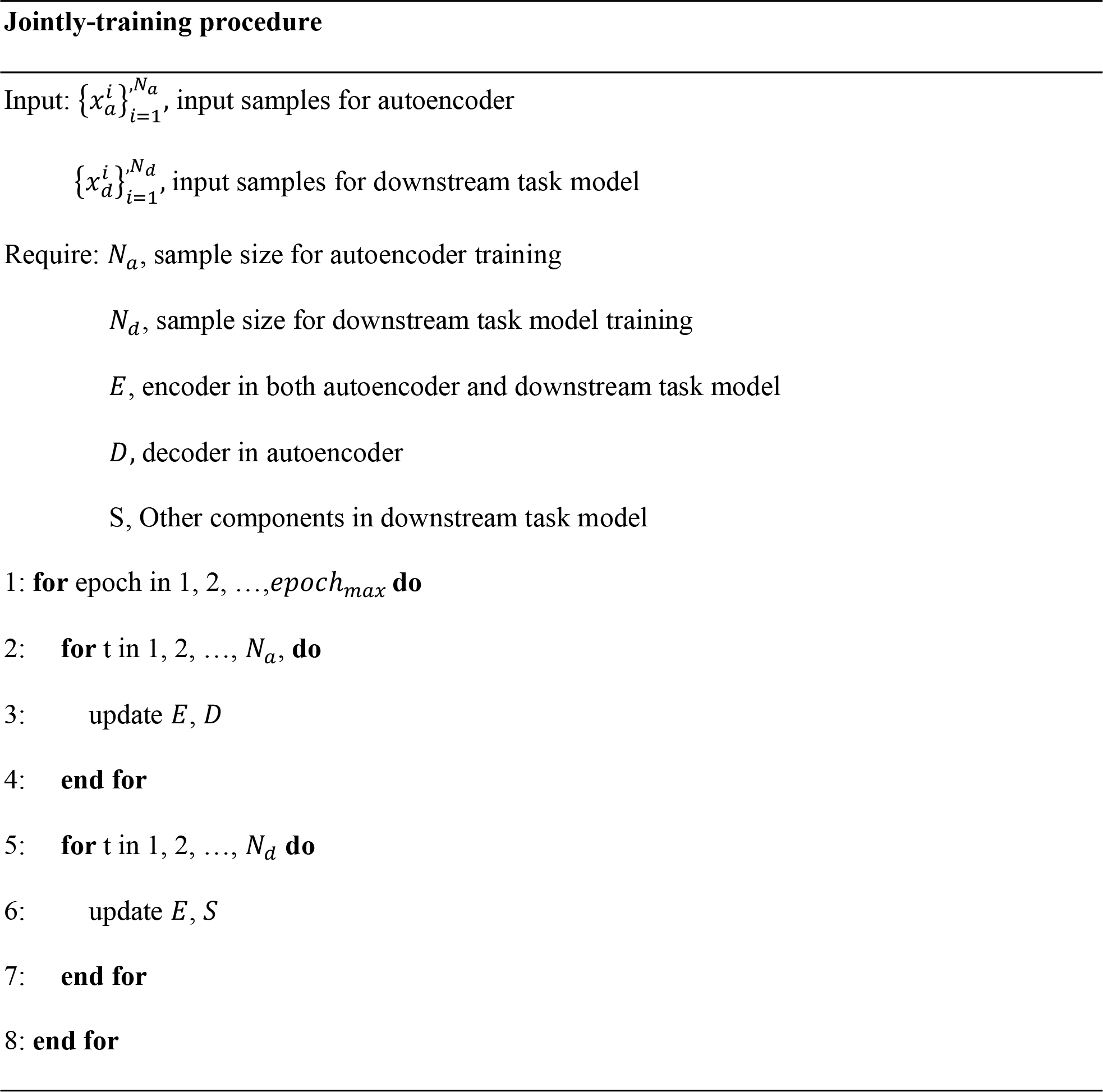

#### Teacher-student model for data augmentation

We augmented our data using a modified procedure described elsewhere^35^. We performed the data augmentation procedure multiple times which is inspired by teacher-student model. In details, we firstly used high confidence data as train data, then we compared our predicted perturbed gene expression to the experimental gene expression profiles for the low confidence data. If the Pearson’s correlations of predicted and experimental profiles are larger than 0.5, we then included this data and the predicted gene expression profile as the next round train data. We eventually performed the data augmentation for 4 rounds (Supplementary Figure 10).

### Model evaluations

#### Chemical transcriptomics prediction

For evaluation purposes, we adopted a leave-new-cells-out cross-validation strategy to split training-dev-test datasets. To be specific, we firstly separated cell lines into different clusters based on their gene expression profile with affinity propagation clustering ^73^(Supplementary Figure 1). We split the training-dev-test dataset to make sure that the cells belonging to one cluster will always be kept in the same fold (Supplementary Table 8). This also means that the cells belonging to different folds will always be dissimilar from each other. During the 3-fold cross-validation stage, we left a group of cells out as the test dataset and the rest as a training-dev dataset, which would be split further to be separate training and dev datasets. Eventually, the performance metrics for these three tests were averaged. This made the problem more challenging than random split. We split the cell lines in the autoencoder carefully so that the cell line in the training dataset for the supervised training will also be in the autoencoder’s training dataset. The same strategies were applied to the dev dataset and test dataset (Supplementary Figure 2). The Pearson’s correlation and Spearman’s correlation between the predicted chemical transcriptomics and ground-truth chemical transcriptomics were used as the evaluation metrics. We also selected the top k upregulated genes and downregulated genes and investigated the model prediction precision for the top-k genes. This precision at k metric could measure the ranking performance of the model.

#### Cell viability prediction

We applied the leave-new-cells-out strategy to evaluate the model performance on the prediction of cell viability. The cells were split into a training dataset, dev dataset, and test dataset. The samples with the same drug were kept in the same fold (Supplementary Table 2). Eventually, the performance from 3-fold cross-validation would be averaged. We used the Pearson’s correlation and Spearman’s correlation scores between the predicted cell viability and ground truth cell viability as the evaluation metrics.

#### Drug side effect prediction

We used the leave-group-of-drugs-out strategy to evaluate the drug side effect prediction performance. For each dataset, we split the data based on the drugs into 5-fold, performed cross-validation, and averaged the performance metrics. Because there are multiple ADR terms to predict, this is a multi-label prediction job. We used micro-ROCAUC, macro-ROCAUC, micro-PRAUC, and macro-PRAUC to evaluate the model’s performance.

### Drug repurposing for Alzheimer Disease (AD)

Gene expression data from brain tissue were downloaded from AD Knowledge Portal. Data from ROSMAP project^74^, MSBB project^75^, and MayoRNAseq^76^ project were uniformly processed by the RNAseq harmonization study into raw count tables. We used all samples from ROSMAP, parahippocampal gyrus samples from MSBB project, and temporal cortex samples from Mayo RNAseq in our study. We then applied MultiDCP framework to predict drug-induced differential gene expression profiles and drug response curves. We trained autoencoder with CCLE, TCGA and AMP-AD data^8, 10^. Our drug candidates include both the approved drugs and drugs under investigation from DrugBank^52^. The tissue gene expression profiles of 46 AD patients and the aforementioned drug candidates were used for the perturbed gene expression profile prediction. We performed personalized differential analysis (PENDA) with the gene expression profiles of AD patient samples and their corresponding control samples^77^. With PENDA, we extracted the up and down-regulated genes for each AD patient and used those genes as their disease signature. 46 patient samples with 10 or more differential expressed genes (among 978 landmark genes) were used for further drug screening. For each patient, we use gene set enrichment analysis (GSEA) to compare their predicted perturbed gene expression profile to their disease signature^53^. Drugs with the lowest negative scores are believed to best reverse the disease signature and are considered as candidate drugs. Then we performed enrichment analysis with the candidate drug category information. The drug category information was extracted from DrugBank^52^. Furthermore, we clustered 46 AD patients into 3 subtypes using mean-shift clustering methods based on their up-regulated and down-regulated gene signatures calculated with PENDA. The up-regulated and down-regulated genes were assigned 1 or -1, and the others are assigned 0. Distance between samples was measured by Manhattan distance. For each subgroup, we integrated the candidate drug category terms and calculated the term frequency.

### Data Availability

The original CCLE, GDSC, TCGA, AMP-AD, L1000, PharmacoDB, SIDER, FARES, and STRING data are publicly available datasets. CCLE data were downloaded from DepMap portal: https://depmap.org/portal/download. GDSC data were downloaded from the GDSC Website: https://www.cancerrxgene.org. TCGA data were downloaded from UCSC Cancer Genome Browser Xena: https://xenabrowser.net/datapages. AMP-AD data were available in https://adknowledgeportal.synapse.org. L1000 data were available in https://github.com/njpipeorgan/L1000-bayesian and https://lincsproject.org. PharmacoDB data could be downloaded from https://pharmacodb.pmgenomics.ca. SIDER and FARES were downloaded from http://maayanlab.net/SEP-L1000/#download. STRING were downloaded from https://string-db.org.

### Code Availability

The code of MultiDCP and data can be accessed through https://github.com/qiaoliuhub/MultiDCP.

## Acknowledgement

This work has been supported by the National Institute of General Medical Sciences of National Institute of Health (R01GM122845) and the National Institute on Aging of the National Institute of Health (R01AD057555). We wish to express our appreciation to Pham, TH., and Zeng, J. from Ohio State University, USA for providing codes of DeepCE: https://github.com/pth1993/DeepCE.

## AD Knowledge Portal

The results published here are in whole or in part based on data obtained from the AD Knowledge Portal (https://adknowledgeportal.synapse.org). First and foremost, the authors would like to acknowledge the patients and their families who provided the invaluable tissue samples to support this work. Rush Alzheimer’s Disease Center, Rush University Medical Center, Chicago provided all ROSMAP data collected, which was funded by; NIA grants: P30AG10161, R01AG15819, R01AG17917, R01AG30146, R01AG36836, U01AG32984, U01AG46152, the Illinois Department of Public Health, and the Translational Genomics Research Institute. Mayo Study data were provided by the following sources: The Mayo Clinic Alzheimer’s Disease Genetic Studies, led by Dr. Nilufer Ertekin Taner (which was not certified by peer review) is the author/funder. It is made available under a CC-BY-NC 4.0 International license. bioRxiv preprint doi: https://doi.org/10.1101/2020.06.29.178590. this version posted June 30, 2020. The copyright holder for this preprint and Dr. Steven G. Younkin, Mayo Clinic, Jacksonville, FL using samples from the Mayo Clinic Study of Aging, the Mayo Clinic Alzheimer’s Disease Research Center, and the Mayo Clinic Brain Bank. Mayo Data collection was funded through; NIA grants: P50 AG016574, R01 AG032990, U01 58AG046139, R01 AG018023, U01 AG006576, U01 AG006786, R01 AG025711, R01 AG017216, R01AG003949, NINDS grant R01 NS080820, CurePSP Foundation, and support from Mayo Foundation.

## The Cancer Genome Atlas Program

The analyses are also part based upon data generated by the TCGA Research Network: https://www.cancer.gov/tcga.

## Author contributions

Q.L. and L.X. designed the computational framework. Q.L. and Y.Q. carried out the implementation and analyzed the data. Q.L., Y.Q. and L.X. wrote the manuscript with input from all authors. L.X. revised the manuscript, conceived and planned the study.

### Competing interests

The authors have declared that no competing interests exist.

## Supplementary information

Supplementary Figure 1. The t-SNE 2D visualization of the cell gene expression profile for all cells after the removal of batch effects. A) All cells in TCGA database are labeled with orange and the cells in the CCLE database are labeled with green. B) The cells are seperated to different clusters based with affinity propagation algorithm. Each cluster of cells are labeled with one color.

Supplementary Figure 2. The training steps for autoencoder model and MultiDCP model. The setup for autoencoder training is shown on the top panel. All cell line data are split to train, dev and test dataset. In the perturbed gene expression profile training stage, the encoder parameters are shared (red arrow). Besides, the cell lines in autoencoder’s training dataset are kept in the training dataset in the MultiDCP training stage (brown). The same can be held for test (yellow) and dev (green) dataset.

Supplementary Figure 3. Predicted drug response curve for selected anti-inflammatory related drug and AD patient tissue pairs.

Supplementary Figure 4. Predicted drug response curve for first 20 selected neural system functions related drug and AD patient tissue pairs.

Supplementary Figure 5. Predicted drug response curve for the remaining 21 selected neural system functions related drug and AD patient tissue pairs.

Supplementary Figure 6. The distribution of log2-transformed gene expression raw counts from different databases, TCGA, CCLE, and AMP-AD. We used the pseudo-count as 0.001.

Supplementary Figure 7. Enriched drug category terms for three AD patient clusters.

Supplementary Figure 8. The statistics of drug response data from different database, CCLE, GDSC1000, CTRPv2, gCSI, and FIMM. The number in each panel is the number of cell lines in each database. The number in the intersection part is the amount of cell lines each database has in common with the CCLE dataset.

Supplementary Figure 9. The distribution of cell viability data before and after we remove the unuseful data in different database, CCLE, GDSC1000, CTRPv2, gCSI, and FIMM. There are some data which has same drug response across the whole dosage range, so we only keep the data in the minimum dosage and maximum dosage.

Supplementary Figure 10. Illustration of Data augmentation procedure.

Supplementary Table 1. Comparison of the MultDC-linear model performance on using gene expression profile and gene dependency profile as the cell line features.

Supplementary Table 2. Hypergeometric test of filtered drug categories for the filter drugs during the AMP-AD drug repurpose study.

Supplementary Table 3. Detailed information about the filtered neural system function related drugs in the AMP-AD drug repurpose study.

Supplementary Table 4. Detailed information about the filtered anti-inflammatory related drugs in the AMP-AD drug repurpose study.

Supplementary Table 5. Enriched functional clusters for three patient groups. The analysis was performed with DAVID.

Supplementary Table 6. Number of cells in training, dev and test dataset for cell viability prediction. We split the cells in three different ways and makes sure the same cell will not appear in different datasets. 886 cell lines are used and are the intersection cell lines set between CCLE and PharmacoDB.

Supplementary Table 7. The number of drugs in training-testing dataset for the 5-fold cross validation used in the drug side effect prediction task. 320 drugs are studied in SIDER dataset and 323 drugs are used in FAERS dataset. Those drugs are found in both these two datasets and the selected low quality LINCS L1000 dataset.

Supplementary Table 8. Number of cell lines in training, dev and test dataset for perturbed gene expression profile prediction task and gene expression autoencoder training task. We evaluated these two tasks with leave new cells out cross-validation. In each split, we leave a group of cell lines out and then split the rest cell lines to training and dev dataset. The cells in training dataset, dev dataset and test dataset are distinct to each other. There are totally 15 cell lines for the perturbed gene expression profile prediction task, which have both gene expression profile found in CCLE and high-quality data in LINCS L1000 project. 11550 cell lines are used for the gene expression autoencoder training and are from both CCLE and TCGA datasets.

## Notes

https://github.com/qiaoliuhub/MultiDCP

### Competing Interest Statement

The authors have declared no competing interest.

