## Supplementary figures and images for "Predictive Modeling of Multiplex Chemical Phenomics for Novel Cells and Patients: Applied to Personalized Alzheimer’s Disease Drug Repurposing"

### Supplementary Figures

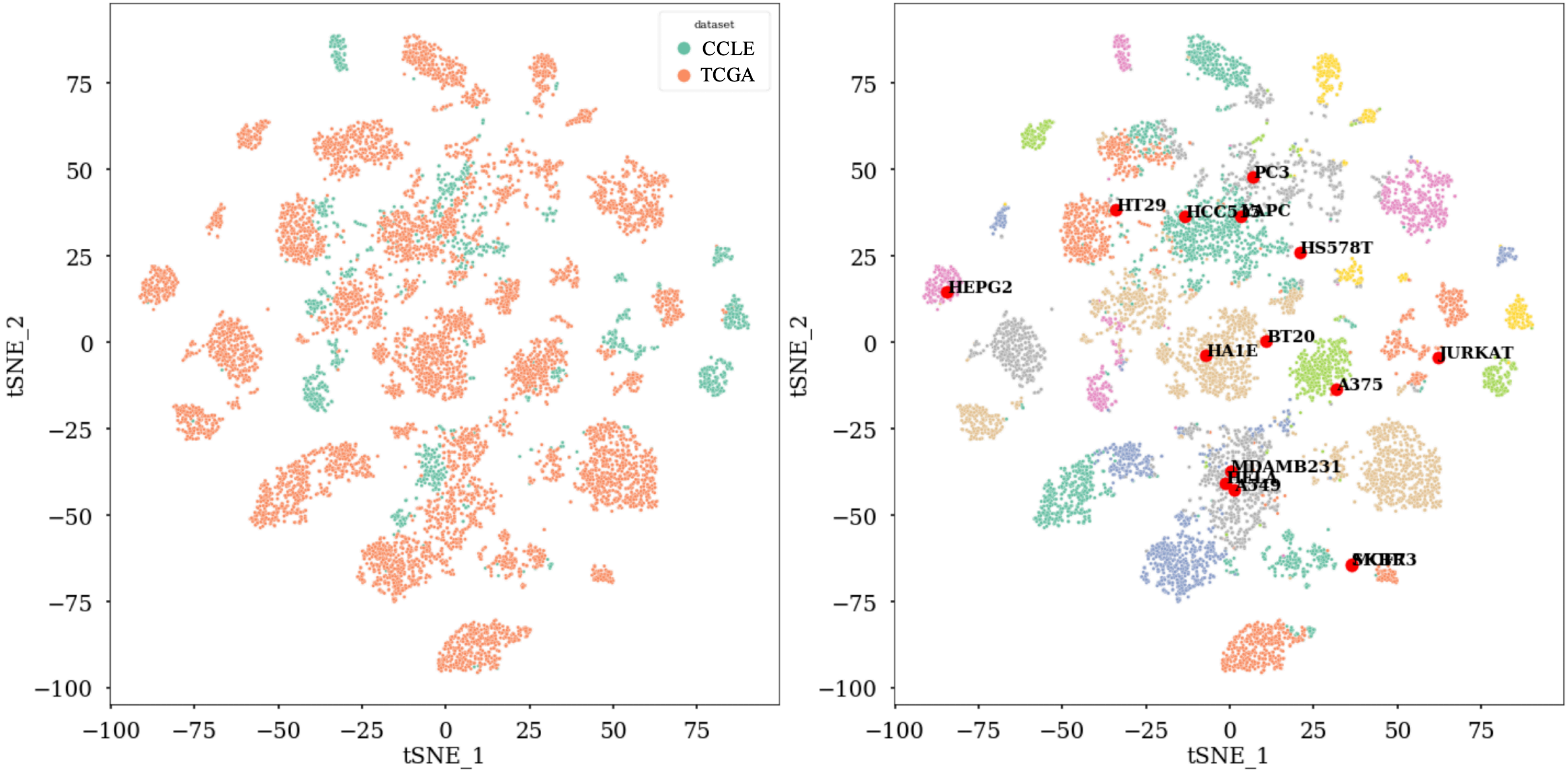

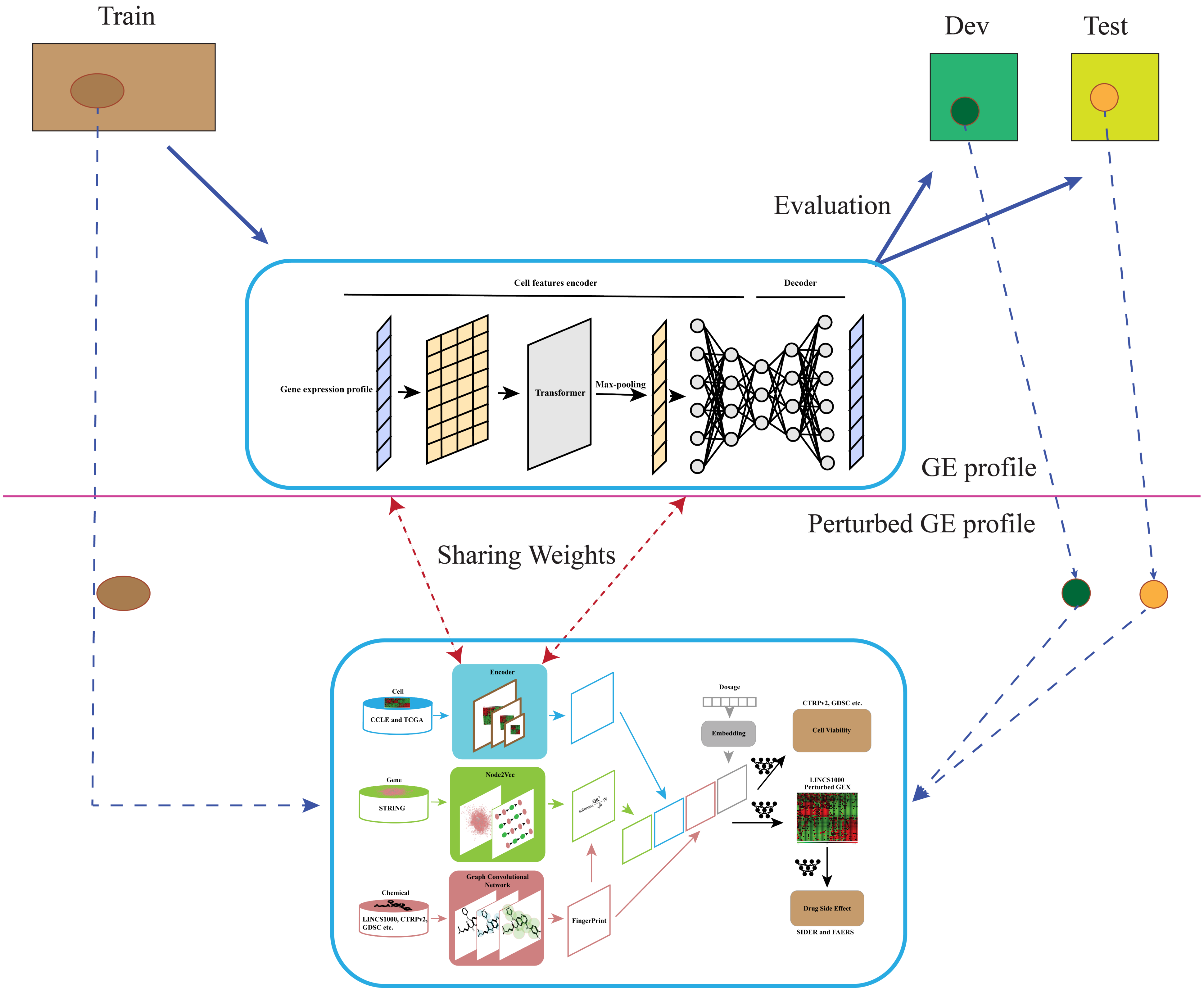

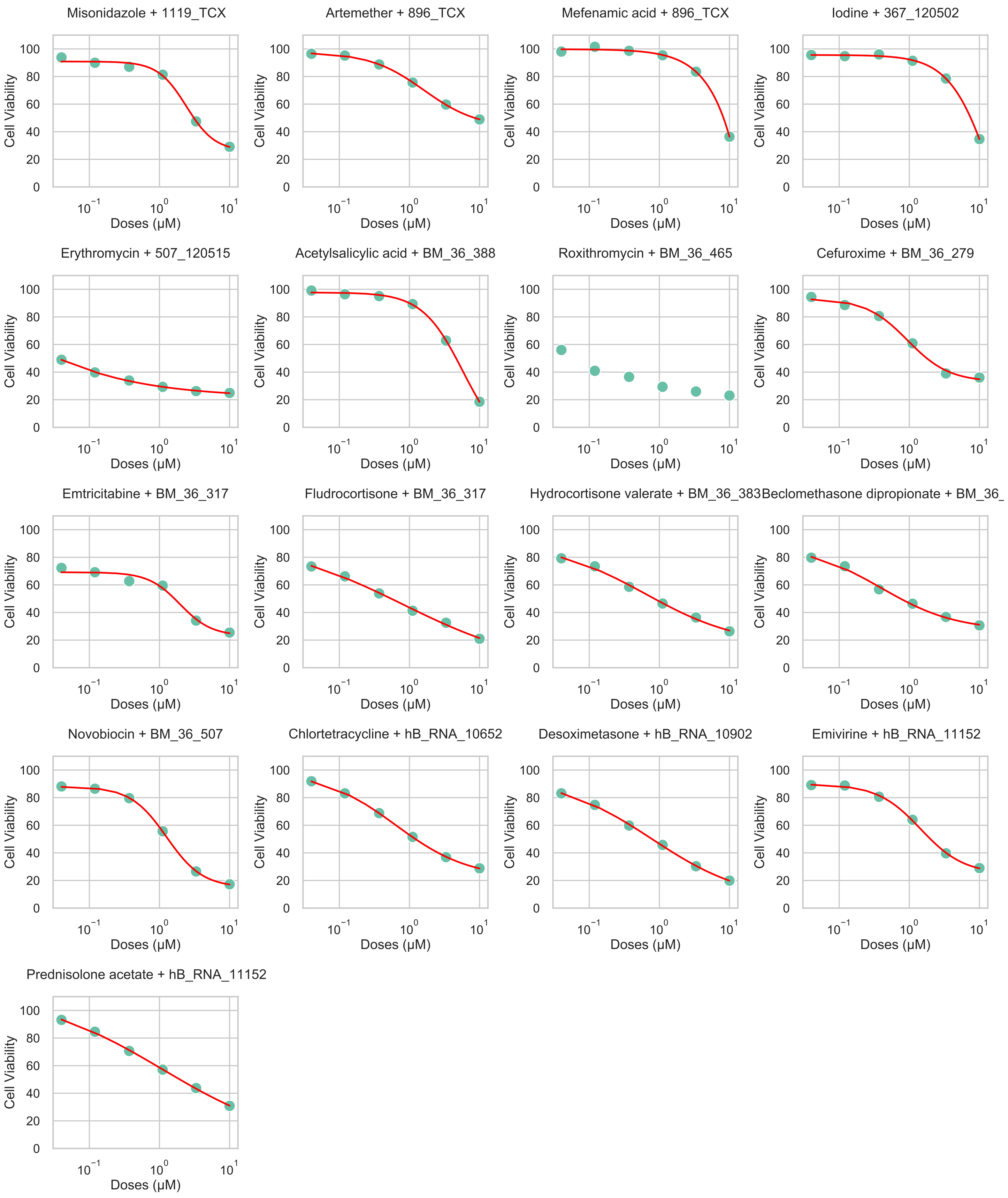

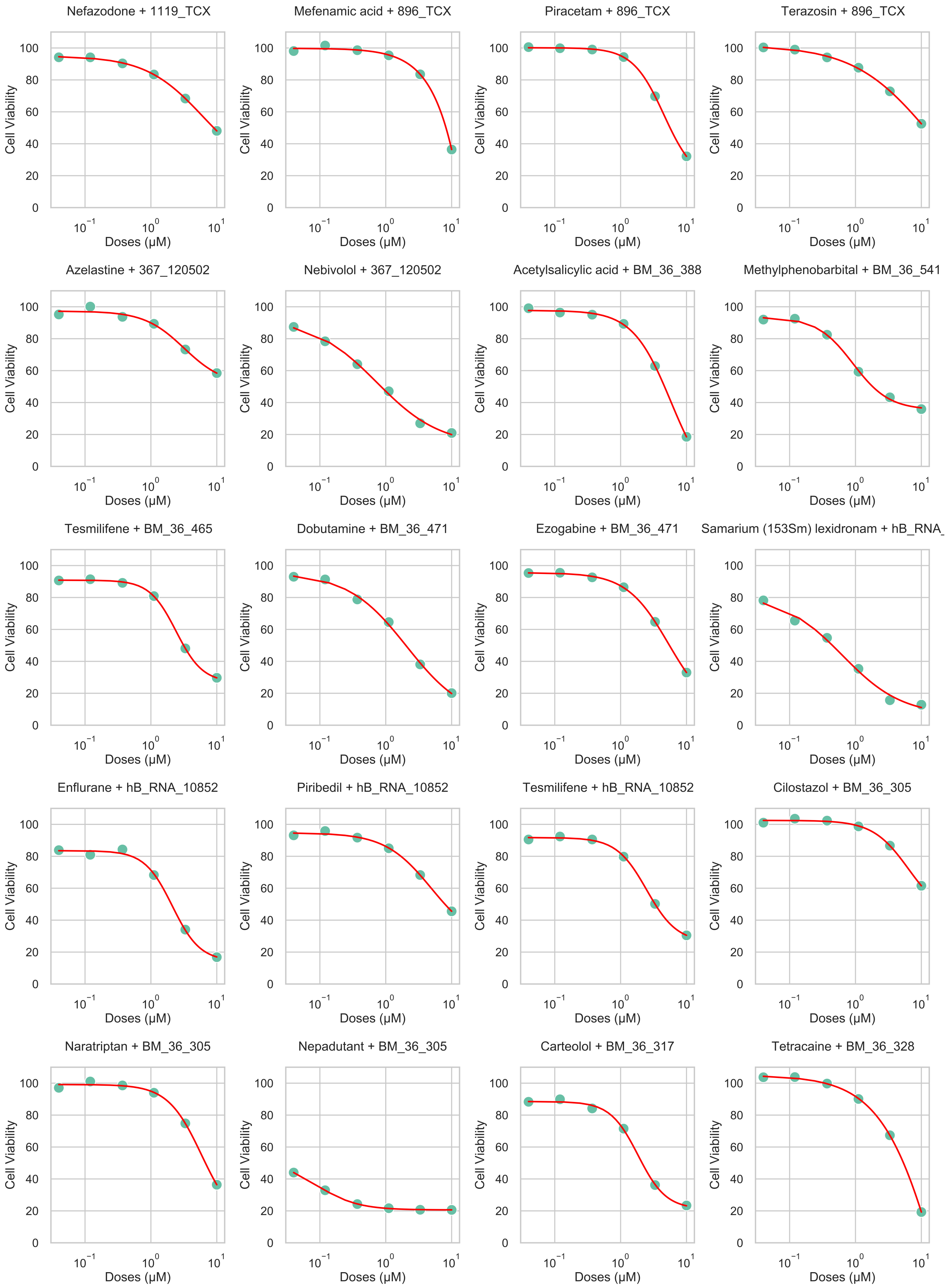

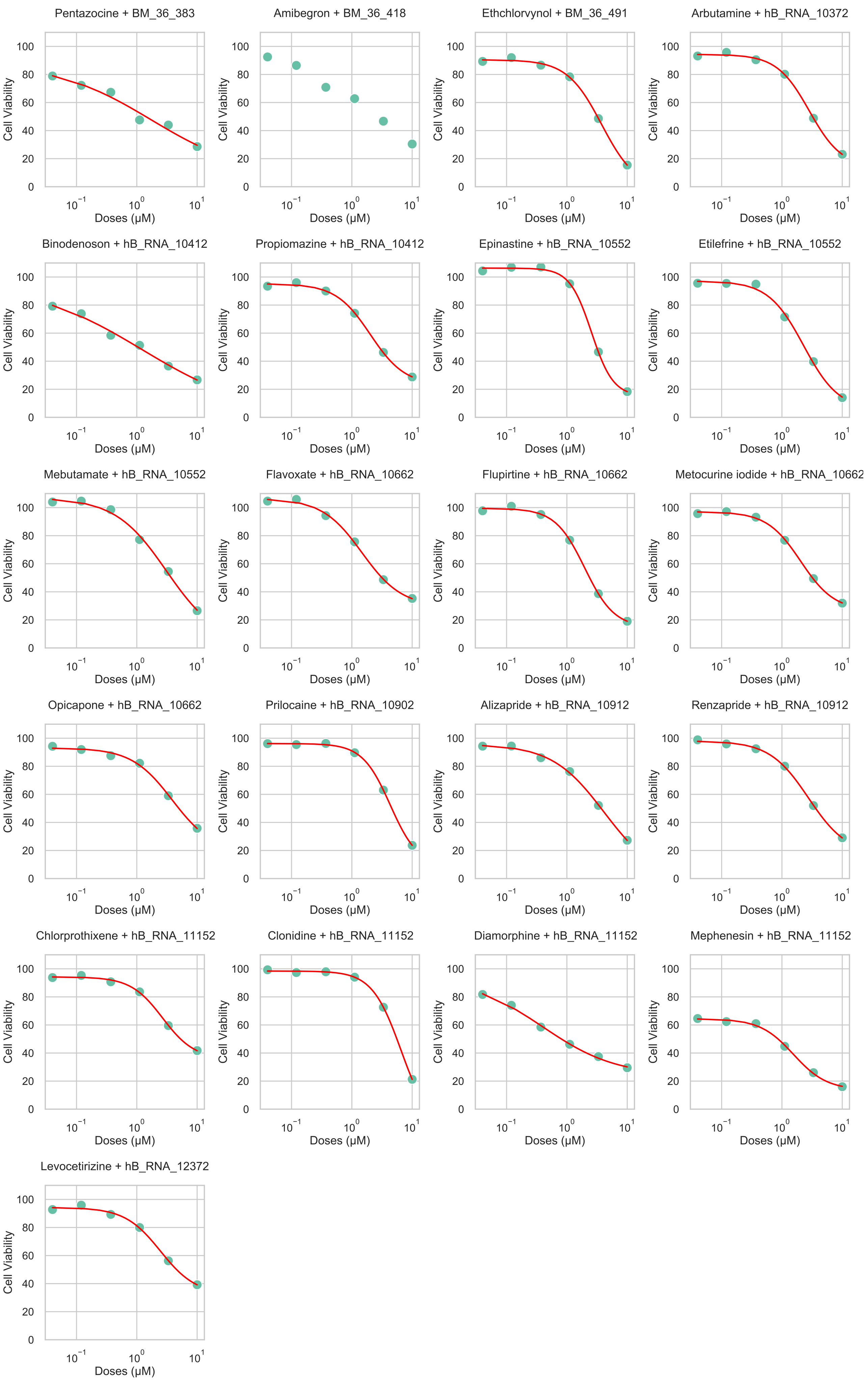

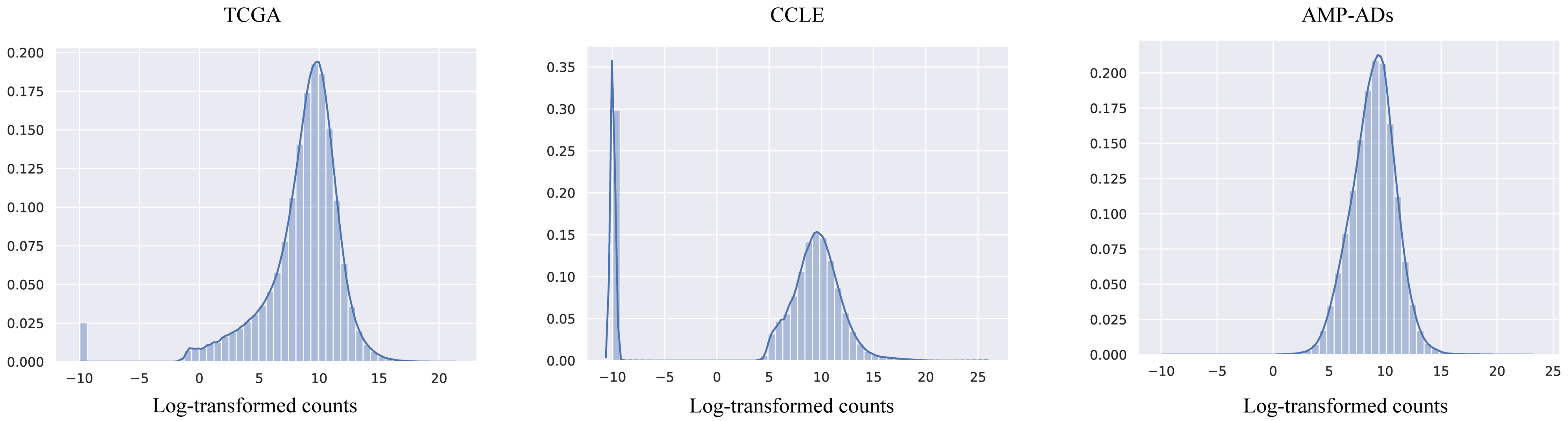

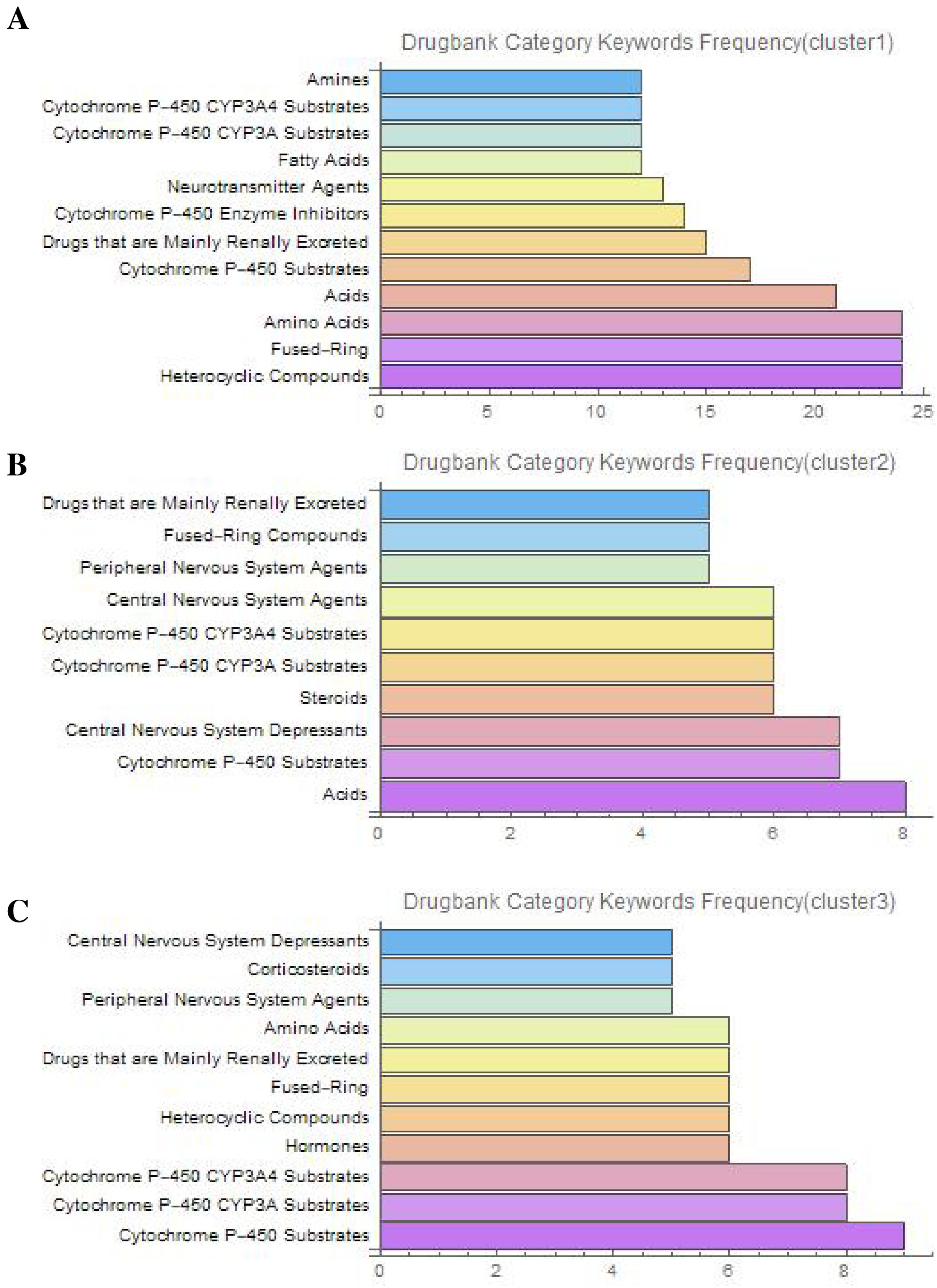

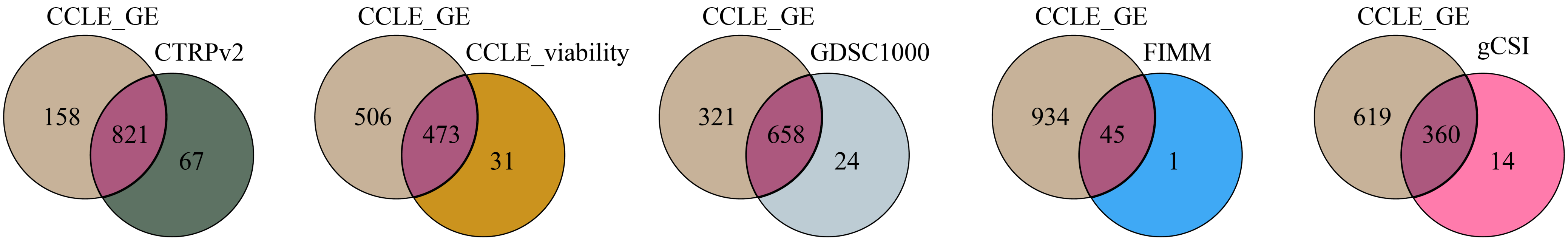

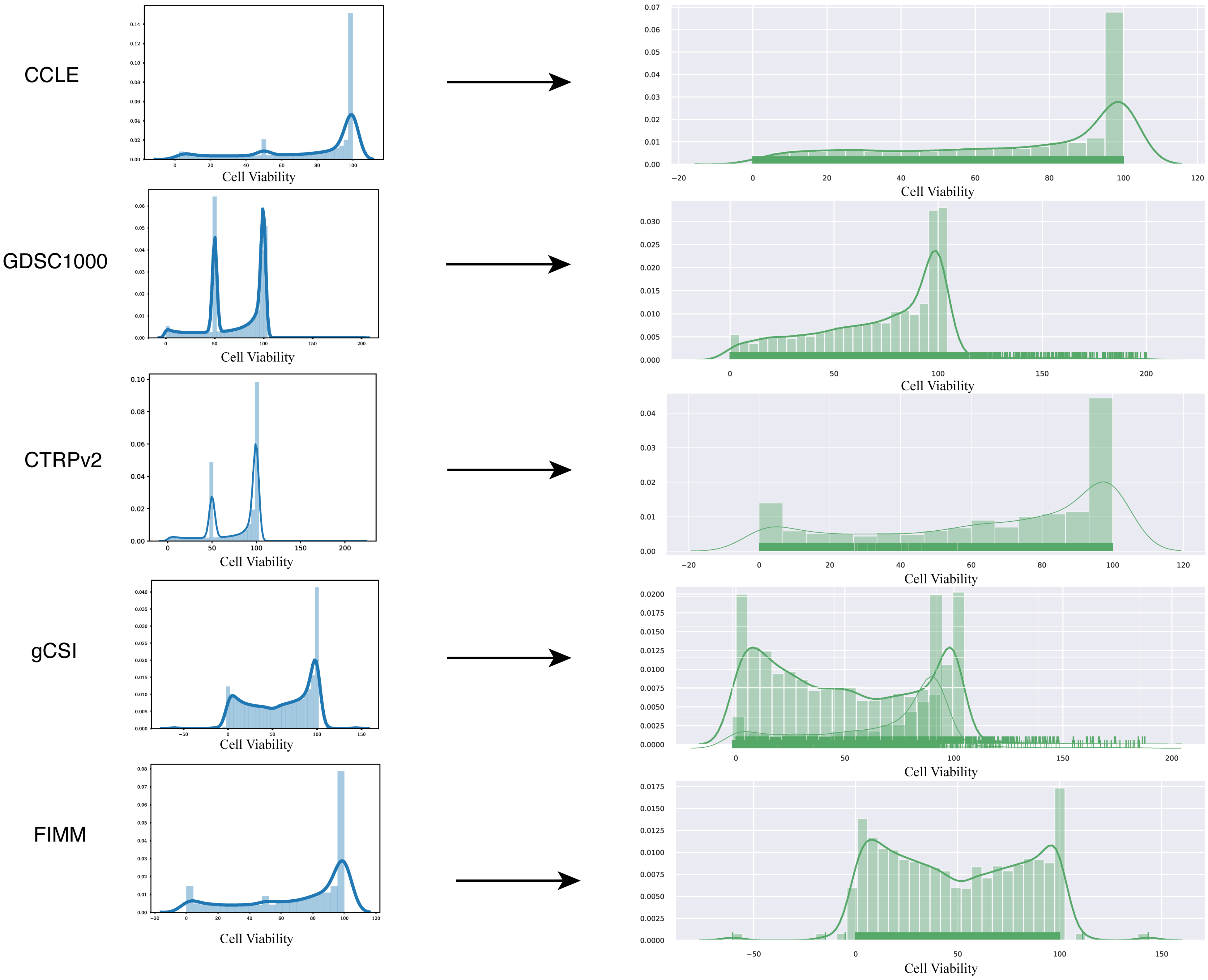

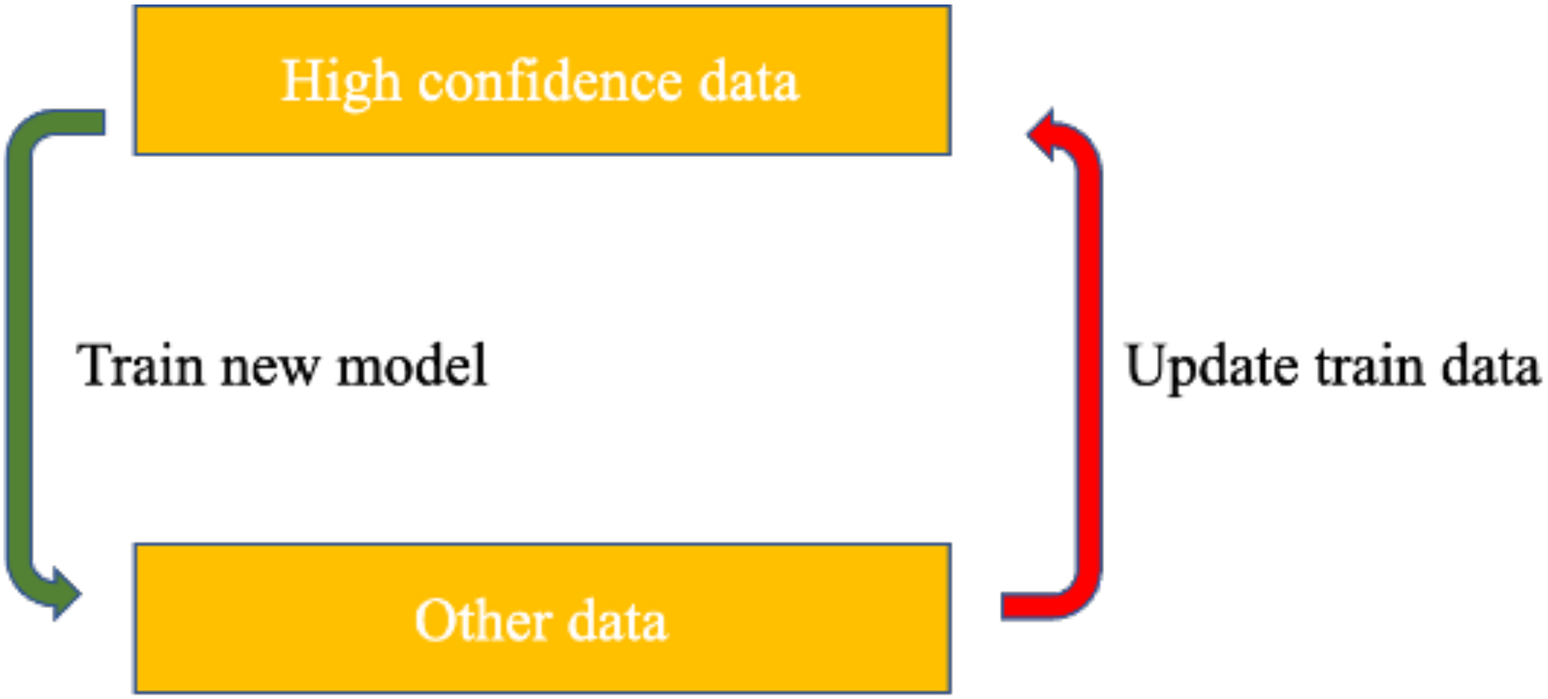
